# Identification of non-charged 7.44 analogs interacting with the NHR2 domain of RUNX1-ETO and exhibiting an improved, selective antiproliferative effect in RUNX-ETO positive cells

**DOI:** 10.1101/2024.06.11.598460

**Authors:** Mohanraj Gopalswamy, David Bickel, Niklas Dienstbier, Jia-Wey Tu, Stephan Schott-Verdugo, Sanil Bhatia, Manuel Etzkorn, Holger Gohlke

## Abstract

The RUNX1/ETO fusion protein is a chimeric transcription factor in acute myeloid leukemia (AML) created by chromosomal translocation t(8;21)(q22;q22). t(8;21) abnormality is associated with 12% of *de novo* AML cases and up to 40% in the AML subtype M2. Previously, we identified the small-molecule inhibitor **7.44**, which specifically interferes with NHR2 domain tetramerization of RUNX1/ETO, restores gene expression down-regulated by RUNX1/ETO, inhibits proliferation, and reduces RUNX1/ETO-related tumor growth in a mouse model. However, despite generally favorable physicochemical, pharmacokinetic, and toxicological properties, **7.44** is negatively charged at physiological pH and was predicted to have low to medium membrane permeability. Here, we identified **M23**, **M27,** and **M10** as non-charged analogs of **7.44** using ligand-based virtual screening, *in vivo* hit identification, biophysical and *in vivo* hit validation, and integrative modeling and ADMET predictions. All three compounds interact with the NHR2 domain and show *K*_D,app_ values of 39-114 µM in Microscale Thermophoresis experiments as well as *IC*_50_ values of 33-77 μM as to cell viability in RUNX1/ETO-positive KASUMI cells, i.e., are ∼5 to 10-fold more potent than **7.44**. **M23** is ∼10-fold more potent than **7.44** in inhibiting cell proliferation of RUNX1/ETO-positive cells. **M23** and **M27** are negligibly protonated or in a ∼1:1 ratio at physiological pH, while **M10** has no (de-)protonatable group. The non-protonated species are predicted to be highly membrane-permeable, along with other favorable pharmacokinetic and toxicological properties. These compounds might serve as lead structures for the optimization of binding affinity, bioavailability, and anti-leukemic effects of compounds inhibiting RUNX1/ETO oncogenic function in t(8;21) AML.

## Introduction

The RUNX1/ETO fusion protein (also called RUNX1/RUNX1T1 or AML1/ETO) is a common chimeric transcription factor in acute myeloid leukemia (AML) created by the chromosomal translocation t(8;21)(q22;q22) (1). This RUNX1/ETO protein inhibits the function of non-modified RUNX1 by blocking myeloid cell differentiation and apoptosis, thereby causing leukemogenesis (2). The t(8;21) abnormality is associated with 12% of *de novo* AML cases and up to 40% in the AML subtype M2 (1,3). Without treatment, AML is a rapidly fatal disease with a median survival time between 10 to 15 months (4). With treatment, AML can be cured in 70% of pediatric patients, 35-40% of adults younger than 60, and 5-15% of adults above 60 years (4,5). However, the 5-year cumulative incidence of relapse is ∼53%, and the overall survival rate is ∼50% (6). Moreover, high-dose chemotherapy is not suitable for elderly patients (7). As a first step towards treating AML after four decades of research, the United States Food & Drug Administration (FDA) has approved ten new small molecule inhibitors that target specific mutations or pathways in the cell cycle (specific AML subsets) (8). However, no targeted treatment of t(8;21)-dependent AML is available at the moment. Being able to target the RUNX1/ETO fusion protein, which is a major driver in t(8;21)-dependent AML, has the potential to enable targeted therapy schemes with superior efficacy and lesser toxicities than conventional chemotherapy (9).

The t(8;21)-generated RUNX1/ETO fusion protein is composed of the DNA-binding Runt-domain encoded by the RUNX1 gene, and four nervy homology regions (NHR1-4) encoded by the ETO gene (10,11). The NHR2 domain (residues 485-552 in RUNX1/ETO) is an oligomerization domain (**Figure S1A**) that plays a role in the dominant detrimental activity of RUNX1/ETO as well as its ability to repress basal transcription (12). Moreover, co-expression of full-length and C-terminally truncated RUNX1/ETO fusion proteins (lacking NHR3-4 domains) causes a substantially earlier onset of leukemia and blocked myeloid differentiation at an earlier stage (13). By contrast, spliced isoforms containing only the NHR1 domain did not exhibit clonogenic potential compared to isoforms containing NHR1 and NHR2 (13,14). Thus, these studies indicated that the tetramerization conferred by the NHR2 domain plays a key role in promoting leukemogenesis (15). Therefore, targeting the oligomerization function of NHR2 emerges as a promising therapeutic strategy for the treatment of AML.

Previously, it has been shown that residues W498, W502, D533, E536, and W540 of the t(8;21)-generated RUNX1/ETO fusion protein, located in the NHR2 domain, are essential for the tetramerization of NHR2 (**Figure S1B**); mutating these ‘hot spot’ residues to alanine resulted in the formation of NHR2-dimers that do not block myeloid differentiation and fail to induce AML in mice (16,17). Thus, targeting these five hot spot residues (**Figure S1C**) by a small molecule inhibitor appears as a promising therapeutic approach to cure AML. An 18-mer peptide containing all hot spot residues interfered with NHR2-mediated oligomerization and could revert the differentiation block (16). The reported inhibition constant (IC), *IC*_50_, for this 18-mer peptide was 250 µM (BS^3^ cross-linking assay) and 390 µM (ELISA) (16).

In general, small-molecule inhibitors are considered to have inherent advantages over peptidic ones, including generally lower production costs as well as better oral availability and membrane permeability (18). Earlier, we identified the small-molecule inhibitor **7.44** (molecular weight 341 Da) (**Figure 1A**), which was shown to (i) specifically interfere with NHR2, (ii) restore gene expression down-regulated by RUNX1/ETO, (iii) inhibit the proliferation, and (iv) reduce the RUNX1/ETO-related tumor growth in a mouse model (19). We subsequently characterized the interaction of **7.44** with NHR2 (affinity *K*_D_ = 4 µM, **Figure S2**) by biophysical experiments and an integrative structural biology approach. In addition, our data showed generally favorable physicochemical, pharmacokinetic, and toxicological properties of **7.44**. However, **7.44** is negatively charged at physiological pH and has been predicted to have a low to medium membrane permeability (20).

**Figure 1.**
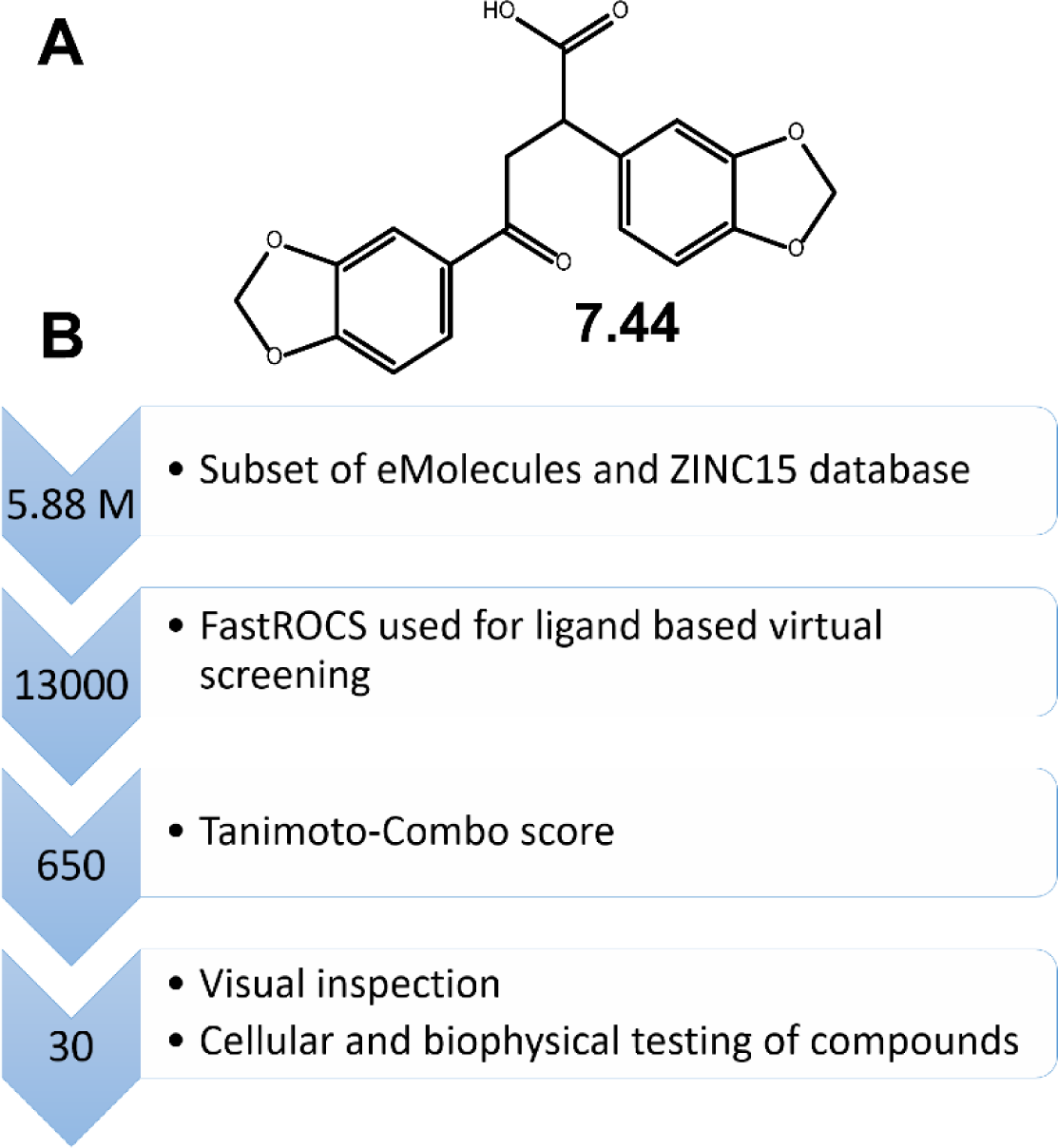
Workflow of ligand-based virtual screening for NHR2 inhibitors. **(A)** Chemical structure of **7.44**. **(B)** FastROCS-based virtual screening using **7.44** as a template was performed on subsets of the eMolecules and ZINC15 databases to obtain 30 compounds for experimental testing.

To obtain a better understanding of the structure-activity relationship (SAR) of inhibiting NHR2 tetramerization, here, we used ligand-based virtual screening with **7.44** as the template scaffold to perform “SAR-by-catalog”. We initially characterized 30 candidates by a cell viability assay in the AML cell line SKNO-1. The three best compounds (**M23**, **M10,** and **M27**), out of the 10 positive hits found by the cell viability assay, interact with the NHR2 domain (STD NMR and nanoDSF measurements) showing *K*_D,app_ values of 39-114 µM (Microscale Thermophoresis (MST)) and *IC*_50_ values of 33-77 μM (cell viability in RUNX1/ETO-positive KASUMI cells). The latter corresponds to a ∼5 to 10-fold increase in potency over **7.44**, likely due to better cell permeability. Likewise, **M23** is ∼10-fold more potent than **7.44** in inhibiting cell proliferation of RUNX1/ETO-positive cells. Integrative modeling further predicted the molecular interactions of **M23**, **M10,** and **M27** with the NHR2 domain. Finally, the three compounds show favorable physicochemical as well as predicted pharmacokinetic and toxicological properties, including a high membrane permeability.

## Results

### Ligand-based virtual screening

To establish a SAR, we performed a ligand-based virtual screening to identify structural analogs of **7.44** using FastROCS (**Figure 1**) (21). FastROCS is an extremely fast shape comparison application, based on the idea that molecules have similar shapes if their volumes overlay well, and any volume mismatch is a measure of dissimilarity. Using **7.44** as a template, the similarity search was done on a subset of the eMolecules and ZINC15 databases containing 5.88 million compounds; the database subsets were chosen based on drug-likeliness. The Tanimoto-Combo score, which considers the molecular shape and pharmacophore according to the automatically assigned standard ROCS color features (donor, acceptor, anion, cation, hydrophobe, rings), was used for ranking the molecules (a higher score indicates more similar molecules, with the maximum being 2). The obtained Tanimoto-Combo scores for the screening results are: 10 molecules > 1.5, 33 molecules > 1.4, 360 molecules > 1.3, 3,800 molecules > 1.2, and 12,500 molecules > 1.0. These scores indicate to what extent substructures of the compounds overlap with **7.44**. Among the top 13,000 molecules, the 650 most similar analogs showing the best Tanimoto-Combo scores were further inspected visually considering the main pharmacophore points. These are the key molecular recognition elements of **7.44** that target the interface of NHR2 according to Metz et al., i.e., the 1,3-benzodioxole moiety enters into a hydrophobic grove formed by W502, L505, L509, and L523, and the carboxyl group interacts with R527 and R528 of NHR2 (16). Finally, we selected and purchased 30 candidates from commercial suppliers for the experimental testing (**Table S1**). All selected molecules have a molecular weight < 400 Da.

### Screening of compounds by cell viability assay and STD NMR

The 30 molecules were initially screened in a cell viability assay after a 96 h treatment utilizing the CellTiter-Glo® assay in the AML cell line SKNO-1 carrying the t(8;21) translocation that results in RUNX1/ETO. Three compounds yielded cell viability with an IC_50_ value in the range of 60 - 150 µM, two compounds in the range of 150 - 250 µM, and five compounds in the range of 400 -650 µM (**Figure 2A**). Particularly, the compounds **M23**, **M10,** and **M27** were found to have the strongest effects on cell viability with IC_50_ values of 62.14 µM, 108.67 µM, and 143.7 µM, respectively. The remaining compounds yielded IC_50_ values ≥ 1 mM indicating no effect on cell viability.

**Figure 2.**
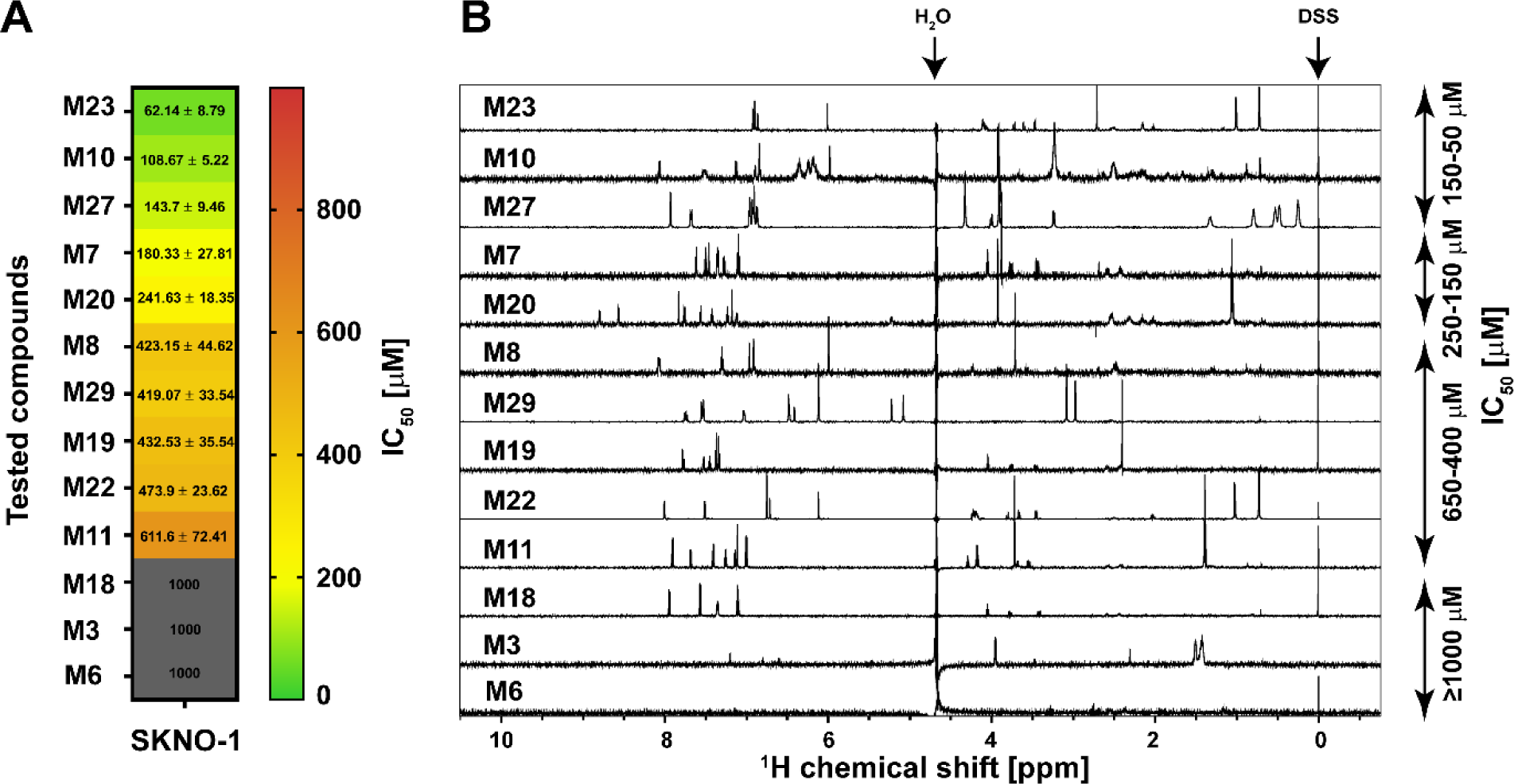
Cell viability assay and STD NMR experiments. **(A)** Anti-proliferative activity of the indicated compounds against the RUNX1/ETO-dependent leukemia cell line SKNO-1. Depicted are the mean ± STD IC50 values in µM calculated by using a sigmoid dose-response curve and nonlinear regression of the raw data normalized to the corresponding DMSO controls. Compounds with IC50 values over 1000 µM were classified as inactive (grey color). Data was collected from three independent experiments (*n* = 3). **(B)** Compounds from the cell-based assay were counter-screened by STD NMR experiments. All compounds with IC50 values ∼50-650 µM (denoted on the right side of the spectrum) showed STD signals in the NMR spectra, indicating their binding to NHR2. **M3** and **M6** showed no STD signals. DSS: Sodium 2,2-dimethyl-2-silapentane-5-sulfonate, used for chemical shift referencing (calibrated to zero ppm).

To test whether compounds identified in the cell viability assay interact with NHR2, saturation transfer difference (STD) NMR spectroscopy was used owing to its high sensitivity and low false-positive rate (22). In the STD NMR experiments, resonances of NHR2 were selectively saturated for 2 s, and the magnetization was transferred to the interacting compound, which resulted in a decreased signal intensity of the bulk ligand. This spectrum was subtracted from a reference spectrum of the same sample recorded in the absence of saturation. Hence, signals in an STD spectrum indicate that a ligand interacts with NHR2 (**Figure 2B**) (23). All 10 compounds with IC_50_ values between 60-650 µM show STD signals. Compounds **M3** and **M6**, which were inactive in the cell viability assay, also did not show STD signals. Hence, **M6** was used as a negative control in the following cell-based assays. **M18** showed STD signals but was not further considered because of the missing efficacy in the cell viability assay.

### Selectivity of hit compounds against RUNX1/ETO-dependent cells

To validate that the efficacy of the three most active compounds in the cell viability assay (**M23**, **M10**, and **M27**) originates from on-target effects at NHR2, the concentration-dependent cell viability in the presence of **M23**, **M10**, or **M27** was measured in RUNX1/ETO-positive KASUMI cells, carrying the t(8;21) gene, and compared to RUNX1/ETO-negative K562 cells, where the NHR2 domain was replaced by the BCR tetramerization domain in the t(8;21) gene. The BCR domain is a coiled-coil tetramerization domain of the oncogenic tyrosine kinase BCR/ABL fusion gene (24) and was chosen because of the high structural similarity, comparable biochemical properties, and because ETO- and BCR-interacting proteins do not show overlap. Yet, the BCR domain lacks the hot spot residues previously identified in the NHR2 domain (17,25) and has been used previously to probe for the selectivity of NHR2-targeting compounds (16).

Compounds **M23**, **M10,** and **M27** reduced the viability of the KASUMI cells with IC_50_ values of 33 µM, 61 µM, and 77 µM, respectively (**Figure 3A** and **Table 1**), indicative of an anti-leukemic effect. **7.44**, measured for comparison, showed an IC_50_ value ∼5 to 10-fold higher (**Table 1**). Compounds **M23**, **M10,** and **M27** were approximately two-fold more potent in KASUMI (RUNX1/ETO^+^) than K562 (RUNX1/ETO^-^) cells, indicating selectivity of the compounds towards RUNX1/ETO-carrying leukemic entities with NHR2 domain. Compound **M6** was used as a negative control and did not show activity in either cell type at a concentration of 1 mM.

**Figure 3.**
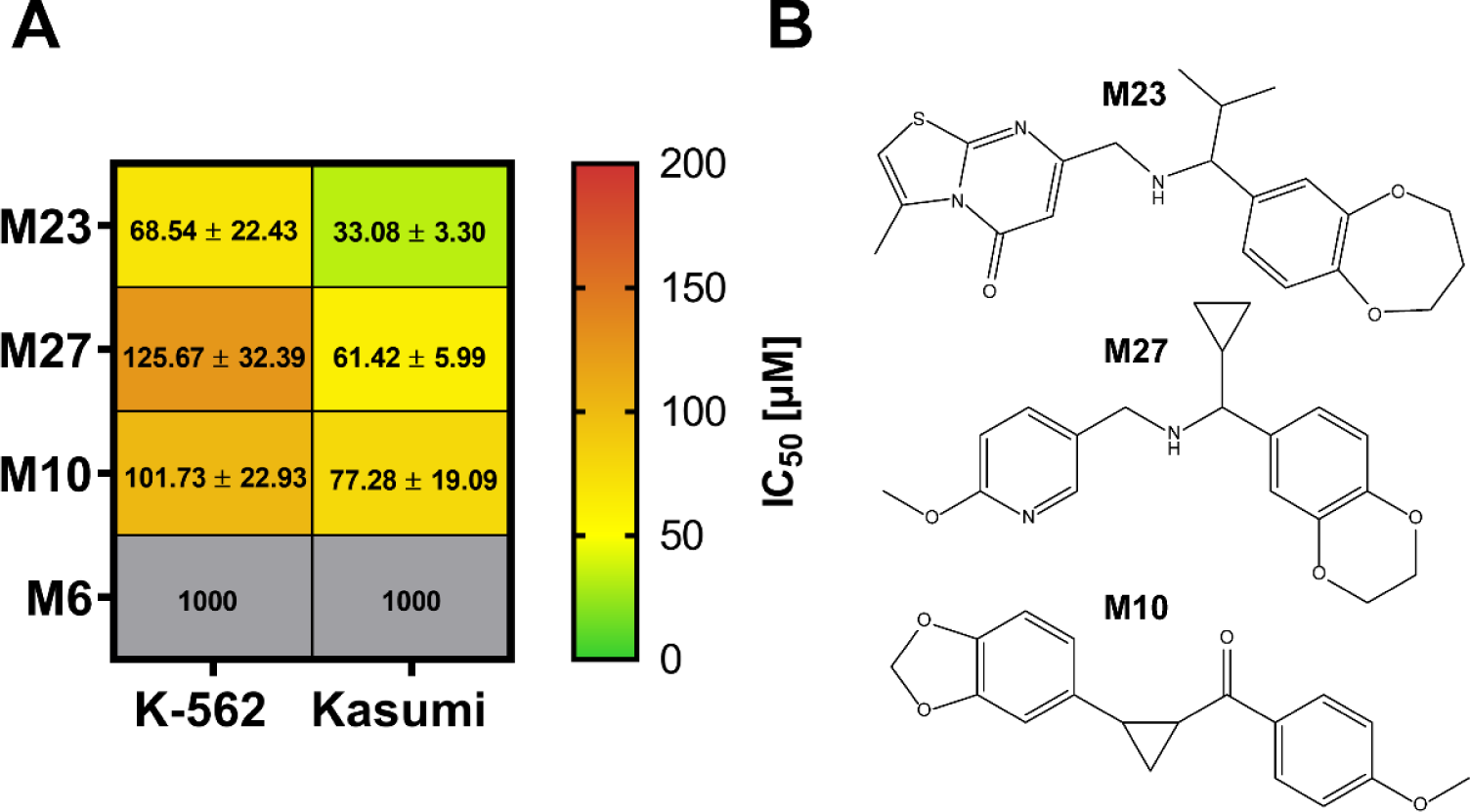
Selectivity of hit compounds against RUNX1/ETO-dependent cells. **(A)** The efficacy of hit compounds **M23**, **M27,** and **M10** identified in the initial screening, as well as the negative control **M6,** was further evaluated in the RUNX1/ETO-positive KASUMI cell line and compared against the RUNX1/ETO-negative K562 cell line. The mean ± STD of the IC50 values are depicted in µM, calculated by using a sigmoid dose-response curve and a nonlinear regression of the raw data normalized to the corresponding DMSO controls. Compounds with IC50 values over 1000 µM were classified as inactive (grey color). Data was collected from three independent experiments (*n* = 3). **(B)** Chemical structures of **M23**, **M27,** and **M10**.

**Table 1.** Anti-leukemic activity and biophysical characterization of lead compounds binding to NHR2. 7.44 was included for comparison.

| Hit compounds | $IC_{50}$ (cell viability) <sup>[a]</sup> | $\Delta T_m$ (nanoDSF) <sup>[b]</sup> | $K_{D,app}$ (MST) <sup>[c]</sup> |
| --- | --- | --- | --- |
| | [μM] | [$^{\circ}$C] | [μM] |
| **M23** | $33 \pm 3$ | $-1.60 \pm 0.03$ | $39 \pm 21$ |
| **M27** | $61 \pm 6$ | $-1.00 \pm 0.09$ | $114 \pm 55$ |
| **M10** | $77 \pm 19$ | $-0.10 \pm 0.04$ | $89 \pm 38$ |
| **7.44** | $357 \pm 16$ | $-0.23 \pm 0.01^{[d]}$ | $2.3 \pm 0.7^{[e]}$ |

<sup>[a]</sup> Viability of RUNX1/ETO-dependent human leukemic KASUMI cells (**Figure 3**); indicated is the mean ± STD ( $n = 3$ ).
<sup>[b]</sup> Shift of the melting point of NHR2 in complex with a 10-fold excess of the compound compared to *apo* NHR2 ( $T_m = 74.5 \pm 0.16$ $^{\circ}$ C) (**Figure 4**); the error comes from the fit ( $n = 3$ ).
<sup>[c]</sup> Apparent $K_D$ of ligand binding to NHR2 (**Figure 4**), obtained by fitting the MST data to a 1:1 binding model; the error comes from the fit ( $n = 2$ ).
<sup>[d]</sup> Compared to previous nanoDSF measurements of **7.44** on NHR2 (20), the **7.44** concentration was now 200 μM.
<sup>[e]</sup> See also **Figure S2**.

### Thermal stability of NHR2 in the presence of hit compounds studied by nano differential scanning fluorimetry (nanoDSF)

nanoDSF was used to assess the thermal stability of NHR2 in the presence and absence of the hit compounds. nanoDSF measures the thermal unfolding transition of a target protein under native conditions, and no extra dye is required (26). Rather, the fluorescence intensity ratio at 350 nm / 330 nm is evaluated, corresponding to tryptophan in nonpolar and polar environments (27). A lower melting temperature *T*_m_ indicates a lower thermal stability (27), as seen previously when **7.44** interacts with NHR2 (20) and confirmed here (**Table 1**).

20 µM of NHR2 was incubated with increasing concentrations of 20, 40, 80, and 200 µM of the respective compound (**M23, M27,** and **M10**), and the melting curves were measured (**Figure S3**). The ratio of fluorescence intensities plotted against the temperature for *apo* NHR2 and in complex with a 1:10 ratio (protein to compound) and the corresponding first-order derivative are shown in **Figure 4A, B**. A melting point of *apo* NHR2 *T*_m_ = 84.1 ± 0.15 °C was observed (**Figure S3A**), which is virtually identical to the one reported earlier (17,20). The addition of 10% DMSO in the buffer containing NHR2 resulted in a destabilization of the protein (*T*_m_ = 74.5 ± 0.16 °C) (**Figure 4A, B**), similar to what has been reported for other proteins in the literature (28). In the presence of 200 μM of the hit compounds, the *T*_m_ of NHR2 decreased by 1.6 ± 0.03 °C for **M23**, 1.0 ± 0.09 °C for **M27**, and 0.1 ± 0.04 °C for **M10** (**Table 1** and **Figure 4A,B**). The *T*_m_ values of the complexes become lower with increasing compound concentrations except for **M10** (**Figure S3).** These results indicate that **M23** and **M27** reduce the thermal stability of NHR2 in a concentration-dependent manner, likely by interfering with the tetramer-dimer equilibrium. It is unclear why **M10** does not show this effect.

**Figure 4.**
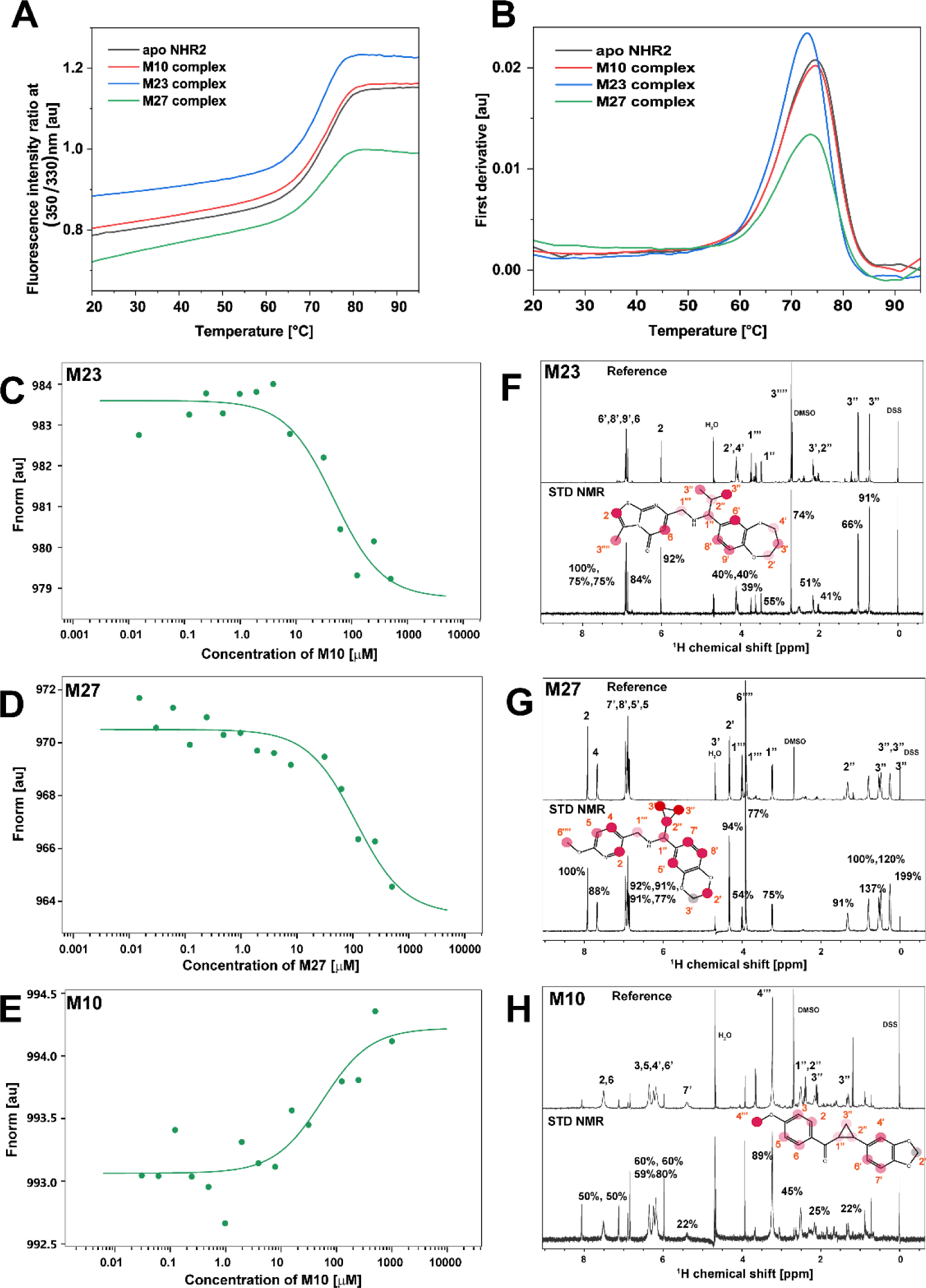
Thermal stability, affinity determination, and ligand epitope mapping of hit compounds M23, M27, and M10. **(A-B)** Thermal unfolding of *apo* NHR2 and in complex with a 10-fold excess of the compounds were studied by nanoDSF (**(A)** Fluorescence intensity ratio at 350 nm / 330 nm; shown are representatives of unfolding events. **(B)** First derivative of the curves in (A)). Changes in the 350 nm / 330 nm fluorescence emission indicate blue- or redshifts. For *apo* NHR2 in 10% DMSO solvent, a *T*m of 74.5 ± 0.16 °C was observed (the error was determined from fitting). Complexes showed reduced *T*m (Δ*T*m of -1.6 ± 0.03 °C for **M23**, Δ*T*m of -1.0 ± 0.09 °C for **M27,** and Δ*T*m of -0.1 ± 0.04 °C for **M10**). **(C-E)** Hit compounds binding to NHR2 detected by MST assay. Titration of hit compounds to a constant concentration of Alexa488 dye-labeled NHR2 induces a change in thermophoresis. The data were fitted to a 1:1 binding model to obtain apparent *K*D values. The inflection point of the curve revealed a *K*D,app of 39 ± 21 µM for **M23 (C)**, 114 ± 55 µM for **M27 (D)**, and 89 ± 38 µM for **M10 (E)**. **(F-H)** Binding of **M23 (F)**, **M27 (G),** and **M10 (H)** to NHR2 studied by STD NMR (lower spectra) (see also Figure 2B) and the corresponding reference 1D (STD-off) ^1^H-NMR (upper spectra). The assignment of the individual peaks is indicated in the chemical structures. The STD effects (Isat / I0) of each proton are indicated on top of the peaks and were mapped as a filled circle onto the structures. Darker red colors denote closer proximity to the protein. The STD effects reveal a particular ligand orientation at the protein.

### Affinity determination by microscale thermophoresis (MST)

MST is a biophysical technique to quantify molecular interactions based on the difference in the movement of molecules along microscopic temperature gradients (29). To determine the apparent dissociation constant (*K*_D,app_) of NHR2 with compounds **M10**, **M23**, and **M27**, 100 nM of dye-labeled NHR2 was mixed with increasing compound concentrations (15 nM to 1 mM). The MST signals were measured for each compound at 24 °C, and normalized fluorescence changes as a function of the compound concentration are shown in **Figure 4C-E**. As the concentration of the respective compound is increased, changes in the thermophoretic curve were observed, implying the binding of the compound to the NHR2 (**Figure S4A-C**). This data was fitted to a 1:1 binding model to obtain *K*_D,app_: 39 ± 21 µM for **M23**, 114 ± 55 µM for **M27**, and 89 ± 38 µM for **M10** (**Table 1**). Previous data of **7.44** (20), reanalyzed for comparison, yielded a *K*_D,app_ of 2.3 ± 0.7 µM (**Table 1**, **Figure S2**), which is within the uncertainty of the dissociation constant of **7.44** binding to NHR2 determined previously (20). The poor solubility of the compounds at higher concentrations gave rise to fluctuations in the data fit and limited reaching a plateau. The *K*_D,app_ values obtained by the MST method are in the same range as the *IC*_50_ values obtained from the cell viability assay except for **7.44** (**Figure 3**, **Table 1**).

### Epitope mapping of the hit compounds by STD-NMR

In addition to identifying molecules that bind to a protein (**Figure 2B**), STD NMR also provides information about the binding epitope of the ligand (22,23). The STD-NMR spectra obtained for NHR2 in the presence of **M23, M27**, or **M10** allow quantifying the STD effect (the ratio of the intensity when the saturation time is 2 s (on-resonance) / the intensity in the reference spectrum (off-resonance), *I*_sat_ / *I*_0_) for each proton (**Figure 4C**). A higher STD effect denotes a closer proximity of the proton to the protein. The STD effects of all three compounds displayed pronounced variations (> 10%), suggesting that the compounds bind to NHR2 through a preferred orientation and not by unspecific binding (30). As an example, the varying STD effects of the isopropyl moiety of **M23** (41%, 66%, 91%) and the cyclopropyl moieties of **M27** (91%, 100%/137%, 120%/199%; values are relative to the peak at 8 ppm) and **M10** (45%, 25%, 22%) are given. The differences in the cyclopropyl moieties might arise from the one of **M27** being more exposed than the one from **M10**, such that the former can interact better. Similar variations were also observed for the methyl groups and aromatic moieties of these compounds. Furthermore, the STD effects of the methyl groups of **M23, M27,** and **M10 (**74%, 77%, 89%) indicate that they are buried upon binding. STD effects of the aromatic protons of **M10** are weaker (∼50-60%) compared to those in **M23** and **M27** (∼75-95%). Overall, the STD data suggests that **M23, M27,** and **M10** do not bind unspecifically to NHR2 and that different structural moieties contribute differentially to the binding. Interestingly, although the *K*_D,app_ values are in the same order of magnitude, the lower STD effects of **M10** parallel the weak effect in nanoDSF of this compound.

### Detection of binding epitopes on NHR2 by molecular dynamics simulations

To study the interactions of **M23**, **M27**, and **M10** with NHR2 on a molecular level, we performed molecular dynamics (MD) simulations of free ligand diffusion (31,32) to simulate the association and dissociation of the compounds to and from NHR2. For **M23** and **M27**, the neutral bases were used as the physiologically most relevant form (see also section “**pK_a_ value determination by NMR”** below). We observed that the compounds interact with NHR2 at multiple sites (**Figure 5** and **Figures S5-S7**). A clustering of the ligand poses based on fingerprints of interactions with NHR2 residues revealed two primary binding epitopes on NHR2. For **M23**, the most populated site (epitope **I**, 13.2%) matches the previously identified binding epitope of **7.44** where interactions with W498 and W502 are formed (20). The second site (epitope **II**, 10.1%) is in the protein-protein interface of the interaction with transcription factor-12 (11). For **M27** and **M10,** the same two epitopes were identified as most populated, although here epitope **II** (15.7 & 13.3%) is more populated than epitope **I** (13.5 & 6.0%). As the occupancies are very similar in the case of **M23** and **M27**, neither one of the binding sites can be identified as preferred.

**Figure 5:**
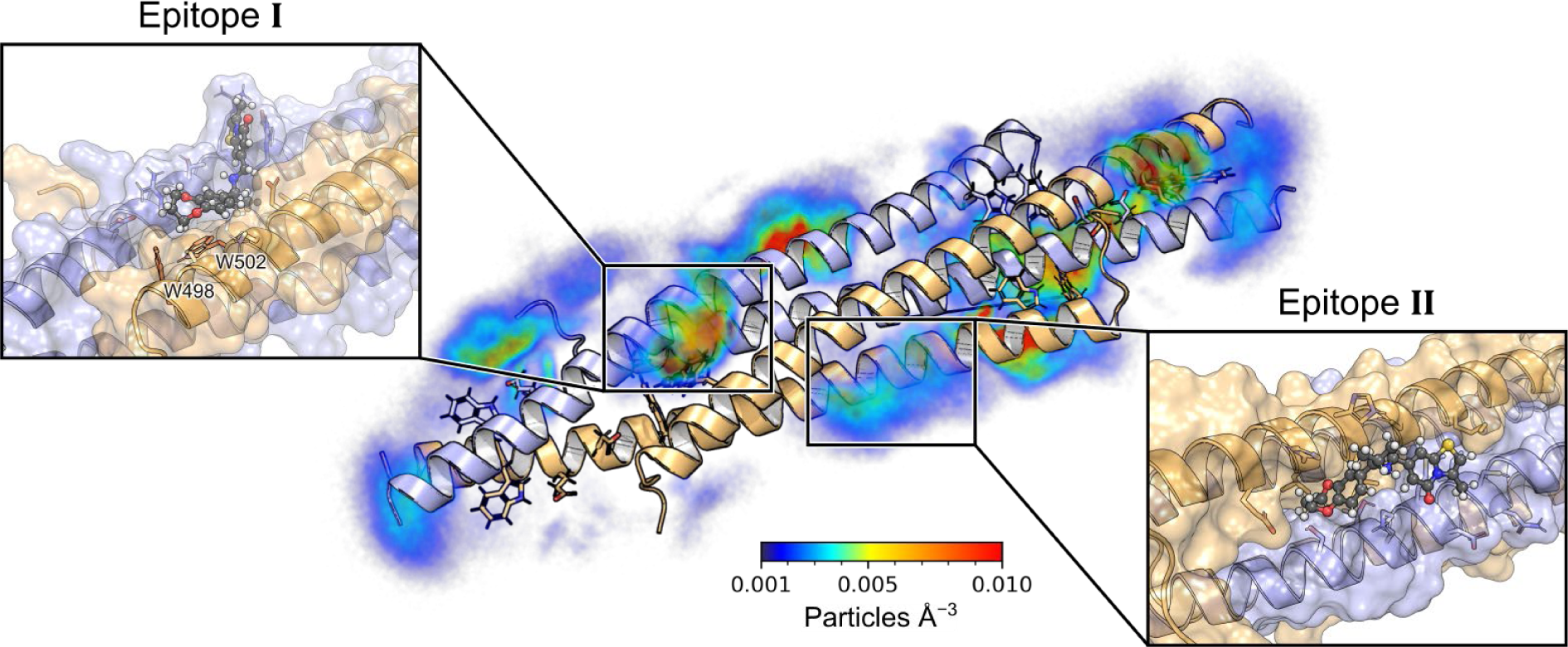
Molecular interactions of compound M23 with NHR2 from MD simulations of free ligand diffusion. On the left side, the heavy-atom particle density of **M23** around the NHR2 tetramer is depicted in a color-coded manner (see color scale). When accounting for the internal symmetry of NHR2, two primary binding epitopes are identified (black rectangles). The left blowup shows **M23** binding to epitope **I**, where the isopropyl group of **M23** is buried between the NHR2 helices, allowing the compound to interact with the hotspot residues W498 and W502. The right blowup shows **M23** bound to the protein-protein interface of NHR2 with transcription factor-12 (11) (epitope **II**). Here, the benzodioxepane ring is partially buried between the helices, while the thiazolopyrimidine ring is bound to a hydrophobic methionine-rich patch.

The STD NMR experiments revealed the isopropyl group of **M23** and the structurally homologous cyclopropyl group of **M27** as the major interacting groups. In the binding mode obtained from epitope **I,** the isopropyl and cyclopropyl groups are buried between the helices of NHR2, which is more predominant for **M23** than **M27**. In contrast, in epitope **II**, both groups are solvent-exposed and only interact transiently with the protein, which is again more predominant for **M23** than **M27**. This might support that epitope **I** is the binding site of **M23** and **M27**. Moreover, the ratio of epitope **I** occupancy over epitope **II** occupancy is largest for **M23** followed by **M27**, which correlates with the relative affinities of these compounds.

### M23 inhibits cell proliferation of AML cells as studied by colony formation assay

A colony formation assay is an *in vitro* cell survival assay based on the ability of a single cell to grow into a colony. We investigated the potency of **M23**, having the lowest IC_50_ and *K*_D,app_ values (Table 1) of the three hit compounds, to inhibit colony formation, which is an important hallmark of myeloid cells (33). For this, a colony-forming unit (CFU) assay was performed in in RUNX1/ETO-positive SKNO-1 and KASUMI cells, and colonies were counted after 14 days (**Figure 6**). **M23** demonstrated a dose-dependent effect on colony formation capabilities and is approximately 10-fold more potent than the previously published NHR2 tetramerization inhibitor **7.44** (19). **M6** was used as a negative control and did not show activity in this assay.

**Figure 6.**
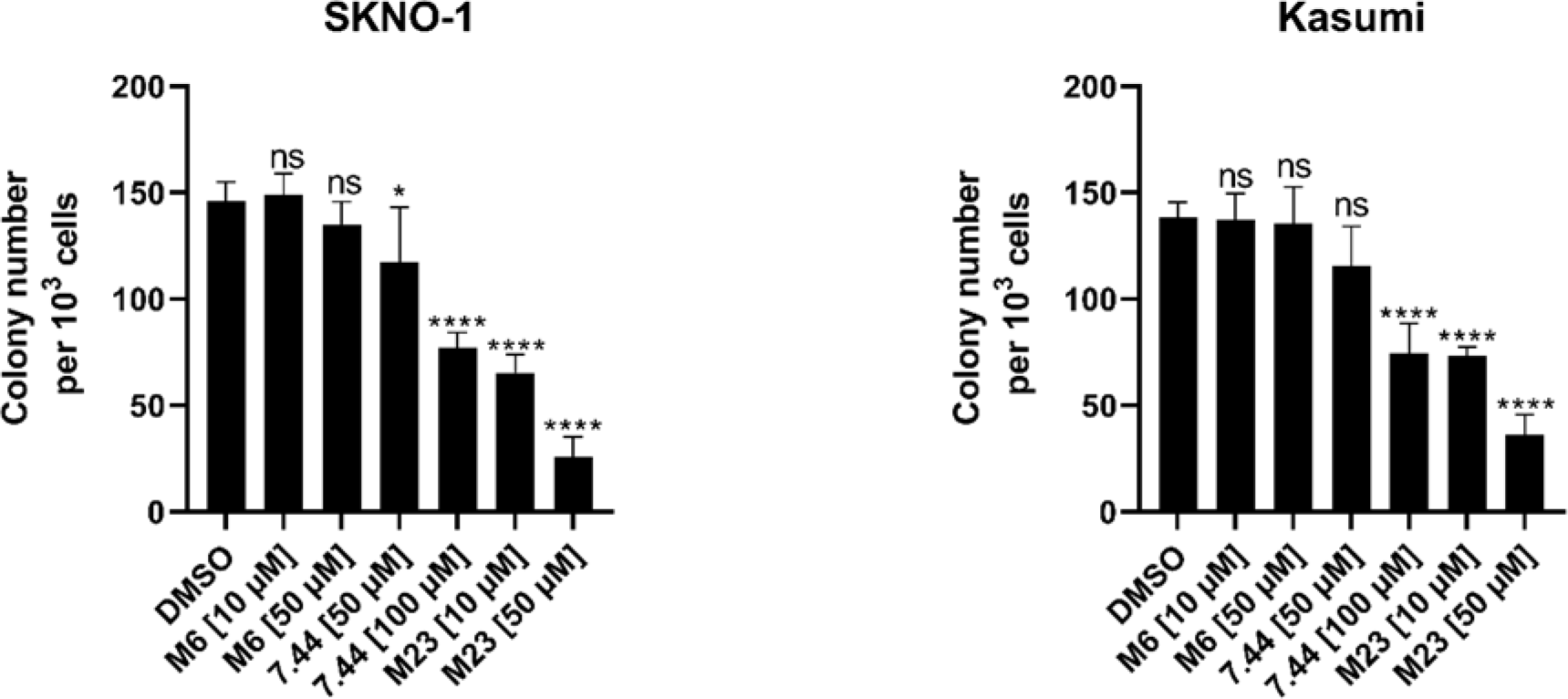
Colony-forming unit (CFU) assay. Representative bar graphs of counted colonies formed by the indicated cell lines 18 days post treatment with **M23**, **M6**, **7.44**, or the control DMSO at indicated concentrations. Statistics was performed using a one-way ANOVA with Dunnett’s multiple comparisons test (\**p* < 0.05; \*\*\*\**p* < 0.0001 vs. DMSO-treated group).

### pK_a_ value determination by NMR

The pK_a_ value is a determinant of physicochemical and pharmacokinetic properties of small molecules. Here, we used NMR titration experiments in the pH range from 2 to 13 to determine the pK_a_ values of **M23** and **M27.** Both compounds contain exchangeable (labile) NH protons (the secondary amines of **M23** and **M27**) that upon (de-)protonation induce different chemical environments for nearby reporter protons (protons 1’’’ and 2’’, **Figure 7, Figure S8**). This results in distinct resonance frequencies of the reporter protons at different pH (**Figure 7A** for **M23**, **Figure S8A** for **M27**); **M10** does not contain a labile proton (**Figure S9**). Sodium 2,2-dimethyl-2-silapentane-5-sulfonate (DSS) was used for chemical shift referencing and calibrated to a zero ppm value in each case. The pK_a_ value was computed from the chemical shift perturbations at the respective pH by fitting to the Henderson-Hasselbalch equation (34) (**Figure 7B, Figure S8B)**. The pK_a_ of **M23** is 6.6 – 6.7, the one of **M27** is ∼7.8 (**Figure 7B, Figure S8B**). These pK_a_ values are ∼4 and ∼3 log units lower than that of dimethylamine (pK_a_ ∼10.7) (35), which results from the electron-withdrawing properties of the nearby aromatic systems. At physiological pH of 7.5, these compounds are thus negligibly protonated (**M23**) or in the ratio ∼1:1 (**M27**).

**Figure 7.**
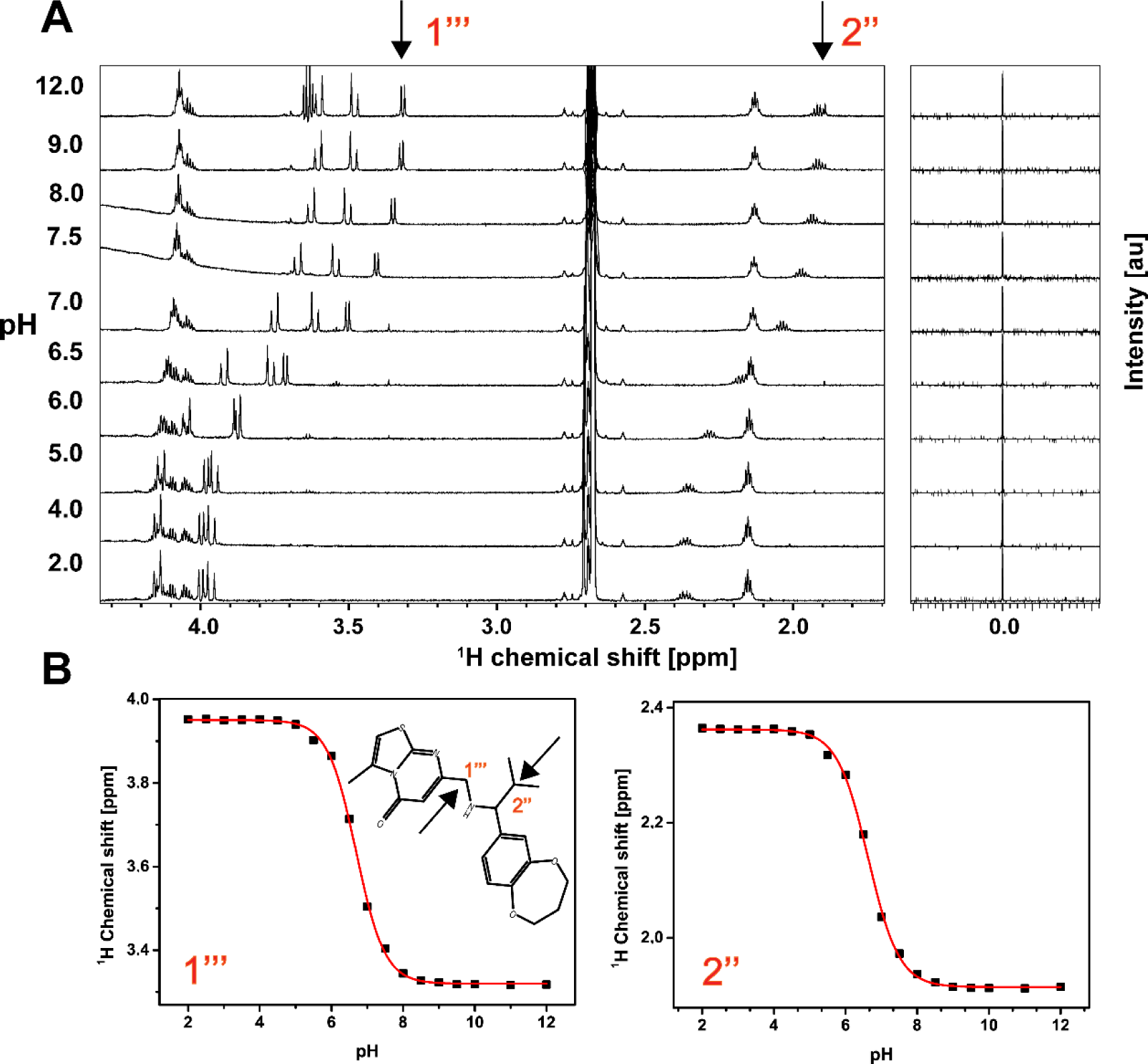
Determination of the pKa value of M23 by NMR. **(A)** Resolved, high-field portion of the 700 MHz ^1^H NMR spectrum of 300 µM **M23** measured in the pH range of 2 to 13 in 50 mM sodium phosphate, 100 mM sodium chloride, 10% (v/v) D2O, 10% (v/v) DMSO-d6. Sodium 2,2-dimethyl-2-silapentane-5-sulfonate (DSS) was used for chemical shift referencing and calibrated to a zero ppm value. **(B)** Chemical shift values of reporter protons (labeled as 1’’’ and 2’’, marked by arrows) were plotted against pH. The pKa value was calculated by fitting to the Henderson-Hasselbalch equation. The pKa value of the secondary amine is 6.69 ± 0.01 and 6.63 ± 0.02 as indicated by protons 1’’’ and 2’’, respectively.

### Membrane permeability prediction of selected compounds

A determining factor for the bioavailability of drugs is their capacity to passively cross cellular membranes (36). Free energy profiles of the permeation process obtained by MD simulations can predict experimentally determined permeabilities from parallel artificial membrane permeability assays (PAMPA), as we previously showed for **7.44** (20,37). Here, we predicted the permeabilities of **M23**, **M27** (both protonated and deprotonated), and **M10** through a PAMPA membrane from MD simulations and configurational free energy computations (**Figure 8A**), considering experimental data from nine reference compounds for calibration (**Figure 8B**, **Table S2**), as done before (see Supporting Information). The computed permeabilities (**Table 2**) classify the species prevalent at physiological pH, non-protonated **M23** and **M10**, as highly permeable (*P*_eff_ > 4.7 x 10^-6^ cm sec^-1^). The same holds for non-protonated **M27**, which exists in a ∼1:1 ratio with protonated **M27**; the latter is low to medium permeable (*P*_eff_ = 7 x 10^-7^ - 2.1 x 10^-6^ cm sec^-1^) (37).

**Figure 8.**
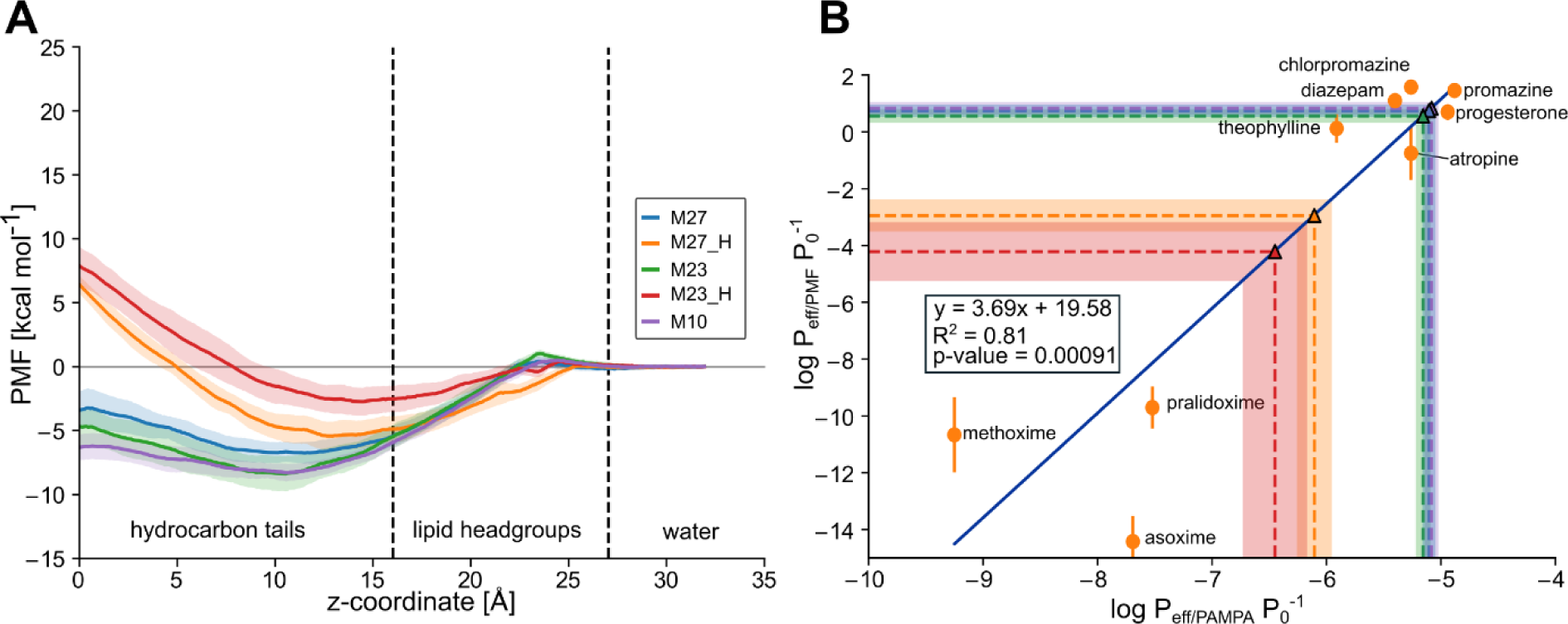
Prediction of membrane permeabilities. **(A)** Potential of mean force (PMF) of permeation of **M23**, **M27**, and **M10**. **M23** and **M27** are additionally considered in their protonated form (**M23_H** and **M27_H**, respectively). As a reaction coordinate, the distance to the center of the DOPC membrane bilayer was used, setting the profile in the water phase to zero. **M10** and deprotonated **M23** and **M27** display favorable free energy in the membrane center, while protonated **M23_H** and **M27_H** display a high energy barrier in the membrane center. **(B)** Calibration curve obtained previously by us (20) between the calculated permeabilities obtained from MD simulations and the experimental values obtained from PAMPA experiments from (37). Using this calibration curve, **M23**, **M27**, and **M10** are predicted to be highly permeable (note the overlap between their values), while **M23_H** and **M27_H** are predicted to be impermeable or low to medium permeable (**Table 2**). *P*eff is the effective permeability, and *P*0 is the unit factor corresponding to 1 cm sec^-1^. Shaded areas correspond to the standard deviation obtained by splitting each calculation into ten slices of 5 ns simulation time.

**Table 2.** Predicted PAMPA permeabilities of compounds M23, M27, and M10.

| Lead Compounds | $\log P_{\text{eff/PMF}} P_0^{-1}$ [a] | $\log P_{\text{eff/PAMPA}} P_0^{-1}$ [b] | $P_{\text{eff}} [\text{cm sec}^{-1}]$ |
| --- | --- | --- | --- |
| **M23** | 0.56±0.23 | -5.16±0.06 | $7.00 \times 10^{-6}$ |
| **M23_H**<sup>[c]</sup> | -4.22±1.04 | -6.45±0.28 | $3.56 \times 10^{-7}$ |
| **M27** | 0.75±0.22 | -5.10±0.06 | $7.90 \times 10^{-6}$ |
| **M27_H**<sup>[c]</sup> | -2.95±0.56 | -6.10±0.15 | $7.86 \times 10^{-7}$ |
| **M10** | 0.83±0.22 | -5.08±0.06 | $8.31 \times 10^{-6}$ |

<sup>[a]</sup> $P_{\text{eff/PMF}}$ , effective permeability in $\text{cm sec}^{-1}$ calculated from free energy calculations. $P_0$ , normalization factor corresponding to $1 \text{ cm sec}^{-1}$ . Errors correspond to the standard deviation obtained from calculating the permeability from ten independent 5 ns simulation slices.
<sup>[b]</sup> $P_{\text{eff/PAMPA}}$ , effective permeability in $\text{cm sec}^{-1}$ calculated from the linear regression obtained from compounds in **Table S2**. $P_0$ , normalization factor corresponding to $1 \text{ cm sec}^{-1}$ .
<sup>[c]</sup> The \_H suffix indicates the molecule is in its protonated (positively charged) state.

### Pharmacokinetic and toxicological property predictions

Pharmacokinetic and toxicological properties were predicted for **M23**, **M27**, and **M10** (**Table 3**). Using Qikprop from the Schrödinger software suite (38), we predicted the central nervous system (CNS) activity, the brain-blood partition coefficient, and the IC_50_ value for blockage of HERG K^+^ channels (**Table 3, No. 1**). All three molecules can be CNS-active, but are predicted to not block HERG K^+^ channels. No violations of Lipinski’s rule of five (39) or Jorgensen’s rule of three (40) are found, judging bioavailability. Furthermore, none of the three compounds has properties falling outside the 95% range of similar values for known drugs. Toxicological properties were predicted using the softwares DEREK Nexus for a variety of endpoints and SARAH Nexus for mutagenicity predictions (41,42) (**Table 3, No. 2)**. No toxicophores were identified in any of the compounds, and the compounds are predicted to be inactive with respect to bacterial mutagenicity.

**Table 3.** Pharmacokinetic and toxicological properties were predicted for M23, M27, and M10.

| No | Description | M23 | M27 | M10 |
| --- | --- | --- | --- | --- |
| 1 <sup>[a]</sup> | CNS | 1 | 1 | 1 |
|  | QplogBB | 0.19 | 0.39 | 0.17 |
|  | QPlogHERG | -6.08 | -6.21 | -4.81 |
|  | Rule of 5 | 0 | 0 | 0 |
|  | Rule of 3 | 0 | 0 | 0 |
|  | #stars | 0 | 0 | 0 |
| 2 <sup>[b]</sup> | Toxicology alert count | 0 | 0 | 0 |
|  | Bacterial mutagenicity | Inactive | Inactive | Inactive |

<sup>[a]</sup> These values were predicted using the Qikprop module of the Schrödinger software suite. CNS: The predicted central nervous system activity (−2 (inactive) to +2 (active)). QplogBB: The predicted brain-blood partition coefficient for oral drugs (recommended range: −3.0 to 1.2). QPlogHERG: The predicted IC<sub>50</sub> value for blockage of HERG K<sup>+</sup> channels (≥ -5 recommended). Rule of 5: number of violations of Lipinski's rule of five (acceptable range: maximum is 4). Rule of 3: number of violations of Jorgensen's rule of three (compounds with fewer (and preferably no) violations). #stars: the number of properties falling outside the 95% range of similar values for known drugs (0 (good) to 5 (bad)) (20).
<sup>[b]</sup> The toxicological properties were predicted with DEREK Nexus and SARAH Nexus. Toxicology alert count: number of predicted toxicophores. Bacterial mutagenicity: predicted bacterial mutagenicity in vitro (20).

## Discussion

We identified **M23**, **M27,** and **M10** as non-charged analogs of **7.44** using a combination of ligand-based virtual screening, *in vivo* hit identification, biophysical and *in vivo* hit validation, as well as integrative modeling and ADMET predictions to characterize interactions with NHR2 or pharmacokinetic and toxicological properties. All three compounds interact with the NHR2 domain and show *K*_D,app_ values of 39-114 µM in Microscale Thermophoresis (MST) experiments as well as *IC*_50_ values of 33-77 μM as to cell viability in RUNX1/ETO-positive KASUMI cells. All three compounds can interact with the previously identified binding epitope of **7.44,** where **M23** is predicted to form interactions with the hot spot residues W498 and W502. NMR-based pK_a_ value determination reveals that **M23** and **M27** are negligibly protonated or in a ∼1:1 ratio at physiological pH, while **M10** has no (de-)protonatable group. The non-protonated species are predicted to be highly membrane-permeable, along with other favorable pharmacokinetic and toxicological properties.

The biophysical characterizations by STD-NMR, nanoDSF, MST, and pK_a_ determination as well as the integrative modeling to determine the binding epitope and predictions of membrane permeability and toxicological properties were performed as recently established or applied by us to characterize **7.44** binding to NHR2 or **7.44** properties (20). For the *in vivo* experiments, RUNX1/ETO-positive or –negative cell lines were used that had been applied before to characterize **7.44** (16,19).

Remarkably, **M23**, **M27,** and **M10** show a ∼5 to 10-fold increase in potency over **7.44** in a cell viability assay with RUNX1/ETO-positive KASUMI cells, although **7.44** is ∼17 to 50-fold more affine to NHR2 according to the *K*_D,app_. **M23** is also ∼10-fold more potent than **7.44** in inhibiting cell proliferation of AML cells. This increased *in vivo* potency can likely be attributed to the prevailing neutral charge of the compounds compared to **7.44**, resulting in improved membrane permeability. Furthermore, **M23** and **M27** are weak bases that might become enriched intracellularly due to a lower intracellular pH (43), which would result in their protonated forms being less membrane permeable. Note that it is important to conserve the low pK_a_ values of the secondary amines for this behavior in subsequent optimization studies. The selectivity factor of ∼2 of the three compounds with respect to RUNX1/ETO-positive over - negative cells is similar to the selectivity found for **7.44** in *in vitro* (16) and *in vivo* (19) assays.

A change in the screening strategy performed now compared to identifying **7.44** can explain the selection of the non-charged **7.44** analogs **M23**, **M27,** and **M10**. Previously, we performed virtual screening focusing on the side chains of hot spot residues in the NHR2 dimer-of-dimers interface as a template, which contained D533, E536, and W540 (16). Unsurprisingly, this resulted in hit suggestions that optimally mimic the hot spot functional groups and frequently contained carboxylic acid(s) or bioisosteres. In addition, we initially evaluated the hit suggestions *in vitro* measuring the inhibition of NHR2 tetramerization, which favored hits with good affinity to NHR2 but neglected the question of availability. By contrast, our virtual screening for analogs now evaluated hit suggestions according to pharmacophore *and* molecular shape similarity, which allowed for the identification of molecules that only sterically mimic the carboxyl group of **7.44** with an isopropyl (**M23**) or cyclopropyl (**M27**) moiety. Next, this time, we first evaluated hit suggestions now using a cell viability assay, which allowed us to assess properties such as cell permeability in addition to a putative action on the target. Only in the second step were the interactions with the target directly probed by STD-NMR. Although we performed only two such steps of complementary virtual screening/hit evaluation campaigns, the positive results suggest that such a strategy could be used to iteratively optimize affinity and pharmacokinetic, in particular, availability, properties.

It may be surprising at first glance why the carboxyl group of **7.44** can be replaced with a neutral carbonyl (**M10**) or even hydrophobic groups (**M23**, **M27**) while keeping the anti-leukemic effect in RUNX1/ETO-dependent cells. Yet, in the predicted binding mode of **7.44** at the NHR2 tetramer, the carboxyl group is oriented towards the solvent whereas the 1,3-benzodioxole moiety inserts in between the helices (20). Thus, an effect-conserving replacement of the carboxyl group is plausible. By contrast, the most active compound **M23** identified here now buries the structurally homologous isopropyl group between the NHR2 helices, allowing the remainder of the compound to interact with hot spots W498 and W502. Although the number of compounds investigated here is too small to derive a quantitative SAR, these findings and qualitative comparison of **7.44** versus **M23**, **M27**, and **M10** suggest that the terminal 1,3-benzodioxole moiety (**M10**), bioisosteres thereof (1,4-benzodioxane (**M27**) or 1,5-benzodioxepine (**M23**)), or even sterically and functionally reduced moieties (methoxyphenyl (**M10**) or methoxypyridyl (**M27**)) together with a distance of three atoms between them are more relevant for interactions with the target. These insights can be exploited in future compound optimizations.

In summary, the identified **M23**, **M27**, and **M10** together with **7.44** might serve as lead structures for further optimization of binding affinity, bioavailability, and anti-leukemic effects of compounds inhibiting RUNX1/ETO oncogenic function in t(8;21) acute myeloid leukemia.

## Materials and methods

### NHR2 protein expression and purification

The expression and purification of the NHR2 domain (residues 485-552 of human RUNX1/ETO fusion protein) was performed as described earlier (20). Briefly, the protein was expressed in *Escherichia coli* Rosetta (DE3)pLysS (Invitrogen) cells that had codon-optimized gene in the pETSUMO vector. The protein was purified by nickel affinity chromatography on a HisTrap column, and the 6His-SUMO tag was removed by incubation with the SUMO protease (1:100 ratio) 50–150 μg/ml followed by affinity chromatography. NHR2 was then purified by S-75 size exclusion chromatography (SEC buffer-50 mM sodium phosphate, 50 mM sodium chloride, 1 mM DTT, pH 8) (GE Healthcare Life Sciences) and analyzed using SDS-PAGE. For NMR experiments, isotope labeling of NHR2 was achieved in ∼98% D_2_O-containing M9 media supplemented with ^13^C-glucose and ^15^N-NH_4_Cl as sole carbon and nitrogen sources, respectively. The labeled protein was purified with the same protocol as the unlabeled protein.

### Inhibitor compounds

All chemical compounds (except **7.44**) were purchased from commercial suppliers (**Table S1**) via MolPort (www.molport.com) and used as received unless otherwise indicated. The compound **7.44** was kindly provided by the Developmental Therapeutics program, National Cancer Institute, USA (NCI code 162496). All compounds were dissolved in DMSO-d_6_ as a 50 or 100 mM stock solution and stored at -20 °C for future use.

### Cell culture

The cell lines K-562 (#ACC 10, BCR-ABL1^+^ CML), SKNO-1 (#ACC 690, RUNX1-RUNXT1 positive AML), and KASUMI (#ACC 220, RUNX1-RUNXT1 positive AML) were purchased from DSMZ (German Collection of Microorganisms and Cell Cultures) and validated by STR DNA profiling. If not stated otherwise, indicated cells were cultured in RPMI 1640 GlutaMax (Gibco, #61870-070) supplemented with 10 % FCS (Sigma-Aldrich, #121031) and 1% Penicillin-Streptomycin (Gibco, #15140-122). SKNO-1 cells were additionally supplemented with 10 ng/ml GM-CSF (Peprotech, #300-03). Cells were kept in a 37°C humidified incubator (BINDER, #BD056) with 5% CO_2_.

### Cell viability screening

Micro-robotic drug library screening was performed as previously described (31). In short, compounds or appropriate controls were printed (randomized) on white 384-well plates (Corning, #3570) spanning eight concentrations in triplicates [1 – 450 μM] in a targeted titration approach utilizing micro-robotics (D300e Digital Dispenser, Tecan). All wells were normalized to the highest vehicle (DMSO) concentration. 30 µL cells were seeded (Multidrop, Thermo Fisher Scientific) with a previously optimized seeding concentration of 3125 cells/ml (K562) or 7000 cells/ml (SKNO-1, KASUMI) in the preprinted plates. After 96 h of incubation (37°C, 5% CO_2_), cell viability was assessed utilizing CellTiter-Glo® luminescence assay (Promega, #G755A) as per manufacturer’s instructions and measured with the SPARK 10M reader (Tecan). The 50% inhibitory concentration (IC_50_) was calculated using a sigmoid dose-response curve and nonlinear regression of the raw data normalized to the corresponding DMSO controls (GraphPadPrism).

### Colony-forming unit (CFU) assay

Assessment of colony-forming units was performed by treating 1 x 10^3^ Kasumi cells with indicated compounds for 48 hours in liquid RPMI1640. Subsequently, treated cells were harvested and plated into 24-well culture plates in triplicates in semisolid MethoCult™ (#04100, STEMCELL) media (33). Colonies were counted after 14 days (31,44).

### Nano differential scanning fluorimetry (nanoDSF) assay

To determine (changes in) the thermal unfolding transition (*T*_m_) of NHR2 in the presence of the compounds, samples were prepared by mixing 20 µM of NHR2, and 20, 40, 80, and 200 µM of respective compounds (**M10**, **M23**, **M27**) to the final volume of 50 µl in a buffer consisting of 50 mM sodium phosphate, 50 mM sodium chloride, pH 8.0, 10% (v/v) DMSO. After incubation for 15 min at room temperature, approximately 10 µl of the sample was loaded into a capillary. The thermal denaturation curves were measured by using intrinsic tryptophan fluorescence of NHR2 (with and without compound) on nanoDSF (NanoTemper, Prometheus NT.40, Germany). The temperature was increased from 20 to 95 °C at a rate of 1 °C/min. The excitation wavelength was 280 nm, and the fluorescence intensity was recorded at both 330 and 350 nm wavelengths (27). The fluorescence intensity ratio F350/F330 was used for the data evaluation (26). For data analysis and fitting to obtain the *T*_m_, the software provided by NanoTemper was used. Duplicates of the same dilution were measured.

### NMR spectroscopy

An STD NMR experiment was performed as previously described (20) at 35 °C on a Bruker Avance III HD spectrometer operating at 750 MHz, equipped with a 5 mm triple resonance TCI (^1^H, ^13^C, ^15^N) cryoprobe and a shielded z-gradient. For STD NMR experiments, 20-30 µM of ^13^C, ^15^N-NHR2 protein were prepared in 20 mM sodium phosphate, 50 mM sodium chloride, 0.5 mM tris(2-carboxyethyl)phosphine, 10% (v/v) DMSO_d6_, pH 6.5, and 10% (v/v) D_2_O. The compound concentration was 1-3 mM in the complex. Selective saturation of protein resonances (on-resonance spectrum) was performed by irradiating at 0.267 ppm for a total saturation time of 2 s. For the reference spectrum (off-resonance), the samples were irradiated at −30 ppm (23). To determine the binding epitope mapping, the STD intensity of the largest STD effect was set to 100% as a reference, and the relative intensities were determined (22,23). Data was acquired with 256 scans, then processed and analyzed by the TopSpin 3.6.1 (Bruker BioSpin) software. Sodium 2,2-dimethyl-2-silapentane-5-sulfonate (DSS) was used for chemical shift referencing.

For pKa value determination, 300 µM of **M10**, **M23,** or **M27** were prepared in 50 mM sodium phosphate, 50 mM sodium chloride, 10% (v/v) DMSO_d6_, 10% (v/v) D_2_O with a pH range of 2-13 (pH of 0.5 steps).1D ^1^H-NMR data was acquired on a Bruker Avance III HD spectrometer operating at 700 MHz spectrometer at 25 °C with 128 scans for each sample. ^1^H chemical shift values of reporter protons were extracted by the TopSpin 3.6.1 (Bruker BioSpin) software and analyzed by the Origin software (OriginLab Corporation, USA). The pKa value was calculated for **M23** and **M27** (**M10** has no labile protons in the tested pH range) compounds by applying the Henderson-Hasselbalch equation as explained by Gift *et al*. (34). DSS was used for chemical shift referencing.

### Microscale thermophoresis

NHR2 labeling was performed with the reactive Alexa Fluor® 488 dye using *N*-hydroxysuccinimide (NHS)–ester chemistry (NHS Ester Protein Labeling kit, ThermoFisher Scientific, USA), which reacts efficiently with primary amines of proteins to form stable dye– protein conjugates. The labeling of NHR2 was performed as described earlier (20). Briefly, 115 µM of NHR2 and 350 µM of Alexa Fluor® 488 dye (410 µl reaction volume) were incubated for one hour at room temperature, followed by 35 min at 30 °C in a buffer consisting of 50 mM sodium phosphate, 50 mM sodium chloride, pH 8.0 (MST buffer). The dye-labeled protein was purified from unreacted free dye using a PD-10 column containing Sephadex G-25 Medium (GE Healthcare Life Sciences) and then centrifuged at 50,000 rpm for 30 min. The concentrations and the efficiency of the labeling were determined as indicated in the manual.

To determine the constant of dissociation (*K*_D_) of the compounds from NHR2, 16 dilution samples were generated by serial dilution (1:1) mixing 200 nM of labeled NHR2 and 1 mM or 2 mM of the respective compound to the final volume of 20 or 25 µl in MST buffer containing 10% (v/v) DMSO. The samples were incubated overnight in the dark before the experiment. The samples were loaded into the Monolith™ NT.115 standard coated capillaries (NanoTemper Technologies), and thermophoresis was measured in the instrument (Monolith NT.115, NanoTemper Technologies) at an ambient temperature of 24 °C, 50% light-emitting diode, and 40% infrared laser (IR) power (MST power) with IR laser on/off times of 30/5 seconds. Triplicates of the same dilution were measured.

The data analysis and the fit (by either thermophoresis with T-jump or thermophoresis or T-jump methods, based on the response amplitude and/or standard error of the MST traces) to obtain the apparent *K*_D_ (considering a 1:1 binding model) was done using the MO affinity analysis software (NanoTemper, Germany) (29). The processed data were plotted with the Origin software (OriginLab Corporation, USA).

### Pharmacokinetic and toxicological prediction

Pharmacokinetic parameters were predicted for the compounds **M23**, **M27,** and **M10** using the QikProp-Schrödinger suite (Release 2023-1) (38). Toxicological parameters were predicted using the toxicology modeling tools DEREK Nexus^®^ (Derek Nexus: 6.0.1, Nexus: 2.2.2) (42) and SARAH Nexus^®^ (Sarah Model—2.0 Sarah Nexus: 3.0) (41) from Lhasa Limited, Leeds, UK.

### Virtual screening

For the virtual screening, the eMolecules (eMolecules Corporate Office, San Diego, CA, USA) and ZINC15 (45) databases were downloaded as SMILES. After removing redundant entries, reactive molecules, fragments, and otherwise unsuitable compounds were filtered out using the OpenEye filter tool with the drug filter. The remaining compounds were protonated at physiological pH using fixpka, and conformers were generated using omega2 (up to 200 conformers per compound). The final dataset was indexed for FastROCS TK (46,47). Using **7.44** as a template (**Figure 1A**), the similarity search was performed on a subset of the eMolecules and ZINC15 databases containing 5.88 million compounds. The Tanimoto-Combo score of each molecule was calculated by considering the shape and pharmacophore according to the automatically assigned standard ROCS color features (donor, acceptor, anion, cation, hydrophobe, rings), and then molecules were ranked. The top 650 most similar analogs showing the best Tanimoto-Combo scores were further inspected visually considering the main pharmacophore points, i.e., if structural elements equivalent to the 1,3-benzodioxole moiety enter into a hydrophobic grove formed by W502, L505, L509, and L523 or equivalent to the carboxyl group interact with R527 and R528 of NHR2 (16). Finally, we selected (molecular weight < 400 Da) and purchased 30 candidates from commercial suppliers via MolPort (www.molport.com) for the experimental testing (**Table S1**).

### Molecular dynamics simulations

#### System preparation

To investigate how the ligands **M10**, **M23**, and **M27** interact with NHR2 on a molecular level, we performed unbiased molecular dynamics (MD) simulations of the diffusion of the ligands around the NHR2 complex. The tetrameric structure of the NHR2 domain (PDB ID 1WQ6 (10), hereafter residue numbering according to the RUNX1-ETO fusion protein (48)) was prepared using Maestro (49); protonation states, Asn/Gln flips, and the tautomeric and flip state of histidines were determined using the built-in implementation of PROPKA. The ligand structures were geometry-optimized in Gaussian 09 at the HF/6-31G* level of theory. For **M27**, the singly deprotonated species was used. Partial charges were assigned according to the restraint electrostatic potential (RESP) fitting procedure (50). For the ligands, GAFF2 (51,52) parameters were used, while ff14SB (53) parameters were used for the protein. For each ligand, initial configurations were built with PACKMOL (54) such that a single ligand molecule was placed at least 20 Å from the central axis of the NHR2 helices. Potassium ions were placed randomly to counter excessive negative charges. Finally, the system was solvated in a truncated octahedron of TIP3P (55) water, leaving at least 11 Å between the edges and any solute molecule. For each system, twenty initial starting configurations were generated.

#### Thermalization and production runs

All MD simulations were performed using the graphics processing unit (GPU) version of pmemd implemented in Amber18 (52,56,57). The Langevin thermostat (58) was used to maintain the temperature at 300 K with a collision frequency of 2 ps^-1^, while the Monte Carlo barostat was used to keep the pressure at 1 bar. The particle mesh Ewald method (59) was used for handling long-range interactions with a cutoff of 8.0 Å. The SHAKE algorithm (60) was used to constrain bonds to hydrogens. Hydrogen mass repartitioning was applied to increase the integration time step to 4 fs for the productive simulations (61).

For thermalization, sequentially water molecules, solute molecules, and, finally, the whole system were minimized for 3000 steps of steepest descent followed by 2000 steps of conjugate gradient algorithm. Positional restraints of 5 kcal mol Å^-2^ were applied to keep other atoms fixed during the minimization. Then, the system was gradually heated to 300 K over 25 ps (NVT), employing a time step of 2 fs, followed by 975 ps (NPT) of density adaptation. The integration timestep was increased to 4 fs, and the system was simulated for further 6,000 ps (NPT), gradually reducing the positional restraints on solute atoms by 0.2 kcal mol Å^-2^ every 600 ns. Finally, the system was simulated for another 2,000 ps (NPT) without any restraints.

After thermalization, all systems (20 starting configurations * 7 ligands) were subjected to 1 μs of unbiased simulation, amounting to a total of 140 μs of MD simulations. Frames were written out at a frequency of 100 ps.

#### Analysis

To identify epitopes on the NHR2 tetramer where a ligand would bind, we calculated the grid densities of ligand heavy atoms mapped onto the tetramer structure using cpptraj (62) from AmberTools of Amber20 (63). For this, we only considered frames where the ligand was in contact (heavy atom-heavy atom distance < 4 Å) with at least five different residues of NHR2 and fitted the NHR2 tetramer structure to a standardized conformer. To characterize the ligand binding, we applied an interaction fingerprint-based hierarchical clustering approach, as described before (20). PyMOL was used for visualization of all molecular data (64).

## Acknowledgements

This work was supported by a grant from the state of North-Rhine Westphalia and the European Fonds for Regional Development EFRE.NRW 2014-2020 to HG. We are grateful for computational support and infrastructure provided by the “Zentrum für Informations- und Medientechnologie” (ZIM) at the Heinrich Heine University Düsseldorf and the computing time provided by the John von Neumann Institute for Computing (NIC) to HG on the supercomputer JUWELS at Jülich Supercomputing Centre (JSC) (user ID: HKF7, VSK33). Financial support by Deutsche Forschungsgemeinschaft (DFG) through funds (INST 208/704-1 FUGG to HG) to purchase the hybrid computer cluster used in this study, GRK 2158/2 (project number 270650915) to HG, and Heisenberg grant ET 103/5-1 to ME is gratefully acknowledged. The authors acknowledge access to the Jülich-Düsseldorf Biomolecular NMR Center. HG is grateful to OpenEye Scientific Software for granting a Free Public Domain Research License.

## Author contributions

MG: Methodology, investigation, formal analysis, writing – original draft; DB: Investigation, formal analysis, writing – review and editing; ND: Investigation, formal analysis; JWT: Investigation, formal analysis; SSV: Investigation, formal analysis, writing – review and editing; SB: Investigation, formal analysis, writing – review and editing; HG: Conceptualization, supervision, project administration, funding acquisition, writing – original draft.

## Data availability statement

All data generated or analyzed during this study are included in this published article (and its Supplementary Information files).

## Additional information

### Supporting information

Supplemental **Figures S1 – S9**.Supplemental **Tables S1-S2**. Supplemental methods for the permeability estimation from molecular dynamics simulations.

