## Supporting Information for "Identification of non-charged 7.44 analogs interacting with the NHR2 domain of RUNX1-ETO and exhibiting an improved, selective antiproliferative effect in RUNX-ETO positive cells"

#### Supplementary figures

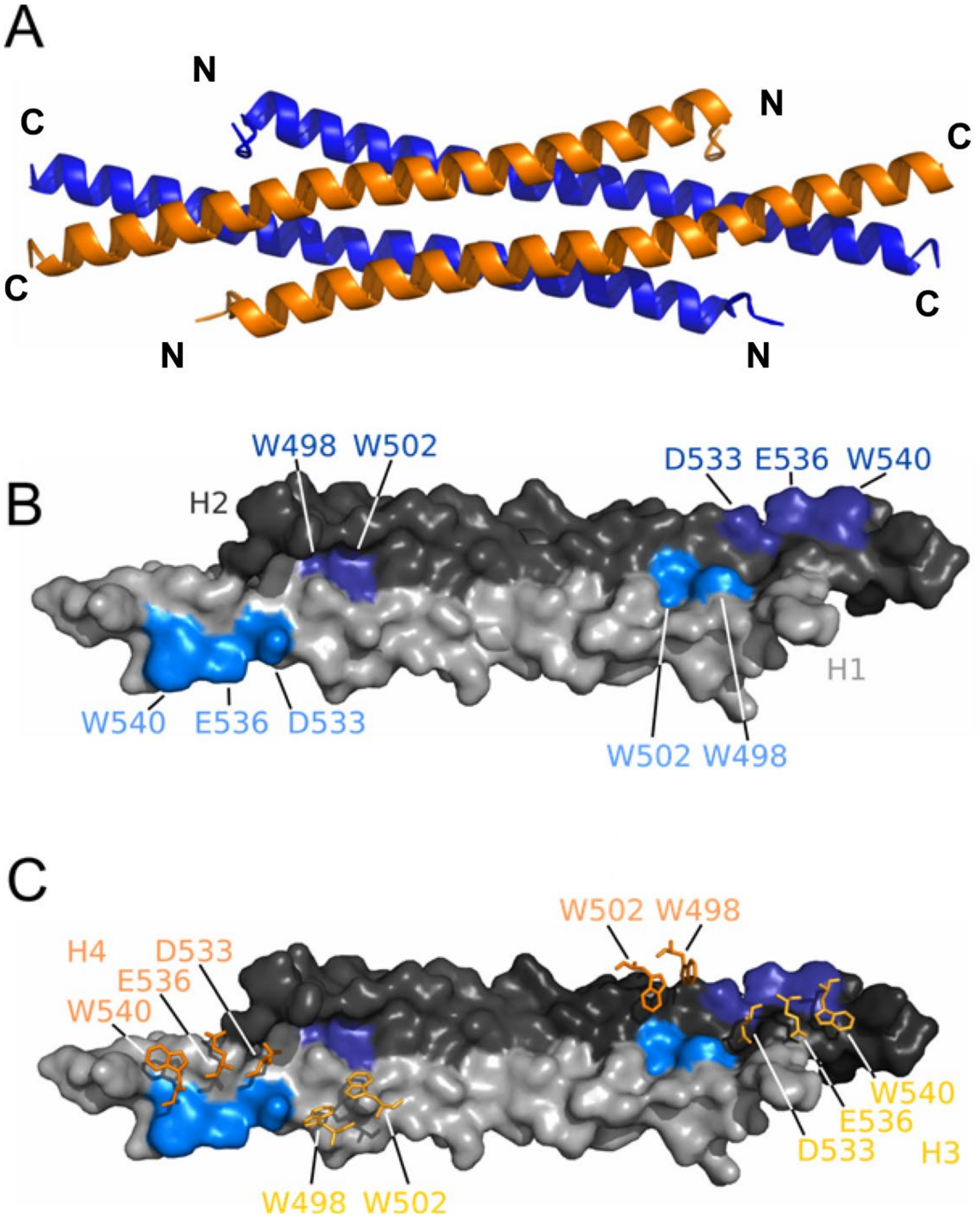


**Figure S1. Structure and hot spots of NHR2**. (A) NHR2 tetramer formed as a dimer of dimers, blue: helix 1 of NHR2 (H1) and helix 2 of NHR2 (H2), orange: helix 3 of NHR2 (H3) and helix 4 of NHR2 (H4). (B) Hot Spots (W498, W502, D533, E536, and W540) of an NHR2 dimer, light grey: surface of H1, dark grey: surface of H2, light blue: hot spots on H1 (surface), dark blue: hot spots on H2 (surface). (C) Representation of the organization of hot spots in the interaction interface of the NHR2 tetramer, in addition to panel C: dark orange: hot spots of H3, light orange: hot spots of H4.

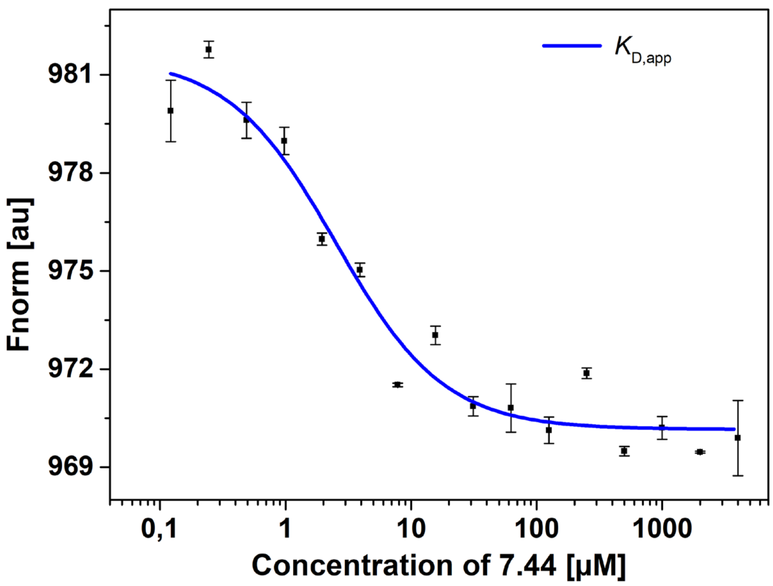


**Figure S2. 7.44 interaction with NHR2 studied by MST assay**. Titration of **7.44** compounds to a constant concentration of Alexa488 dye-labeled NHR2 induces a change in thermophoresis. The data previously obtained in ref. (1) were fitted to a 1:1 binding model to obtain an apparent dissociation constant $K_{D,app}$ of 2.3 ± 0.7 µM (blue).

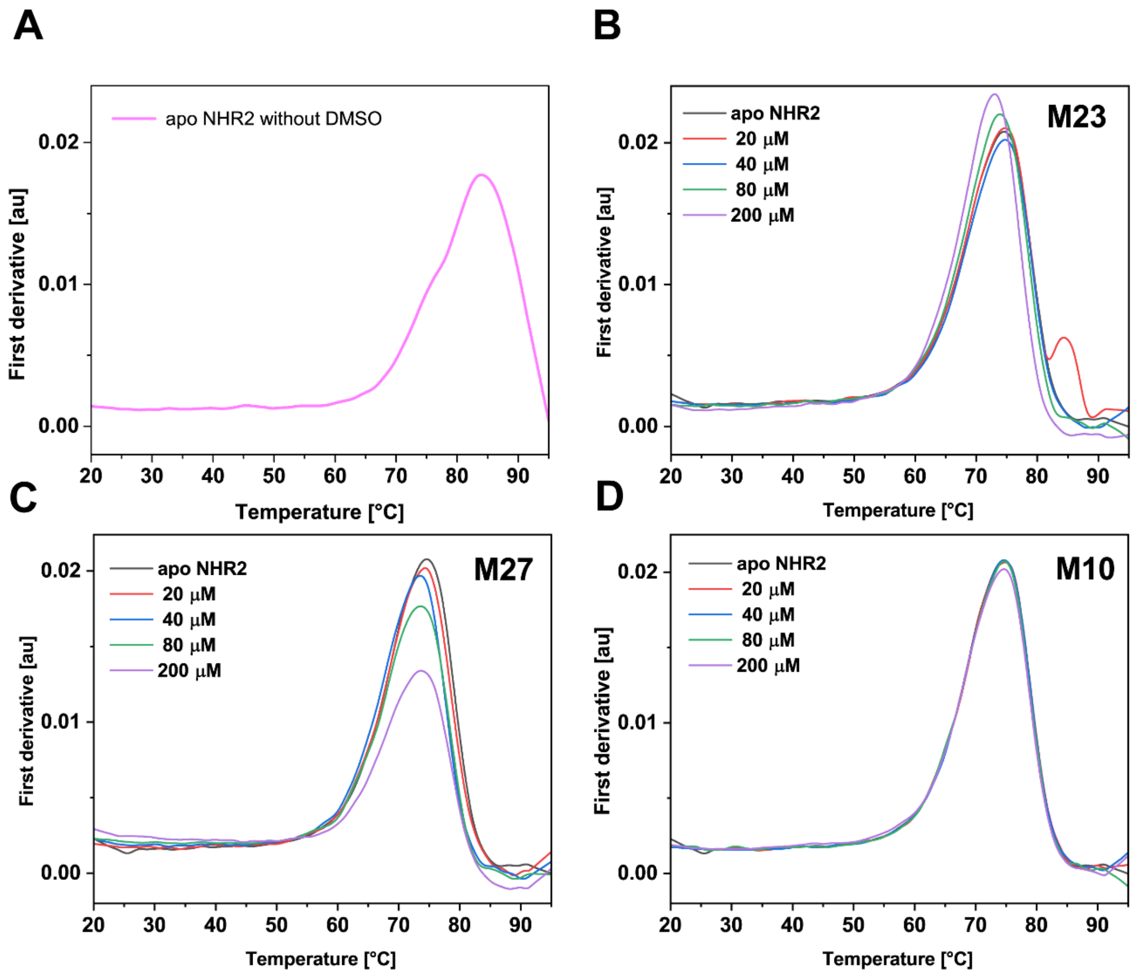


**Figure S3. Concentration-dependent melting curves of NHR2 measured by nanoDSF**. To determine the thermal unfolding transition ($T_m$) of *apo* NHR2 without 10% of DMSO **(A)** and then in the presence of the hit compounds, samples were prepared by mixing 20 µM of NHR2 and 20, 40, 80, and 200 µM of the respective compound **(B) M23**, **(C) M27**, or **(D) M10** to the final volume of 50 µl in a buffer consisting of 50 mM sodium phosphate, 50 mM sodium chloride, pH 8.0, 10% (v/v) DMSO. Changes in the ratio of fluorescence emission at 350 nm / 330 nm indicate blue- or redshifts. The first-order derivatives of the thermal unfolding events are shown. A $T_m$ of 84.1 ± 0.15 °C (panel A) and 74.5 ± 0.16 °C (panels B-D) were observed for *apo* NHR2 without and with 10% of DMSO, and the presence of the compounds lowered the $T_m$ with increasing compound concentration except for **M10**.

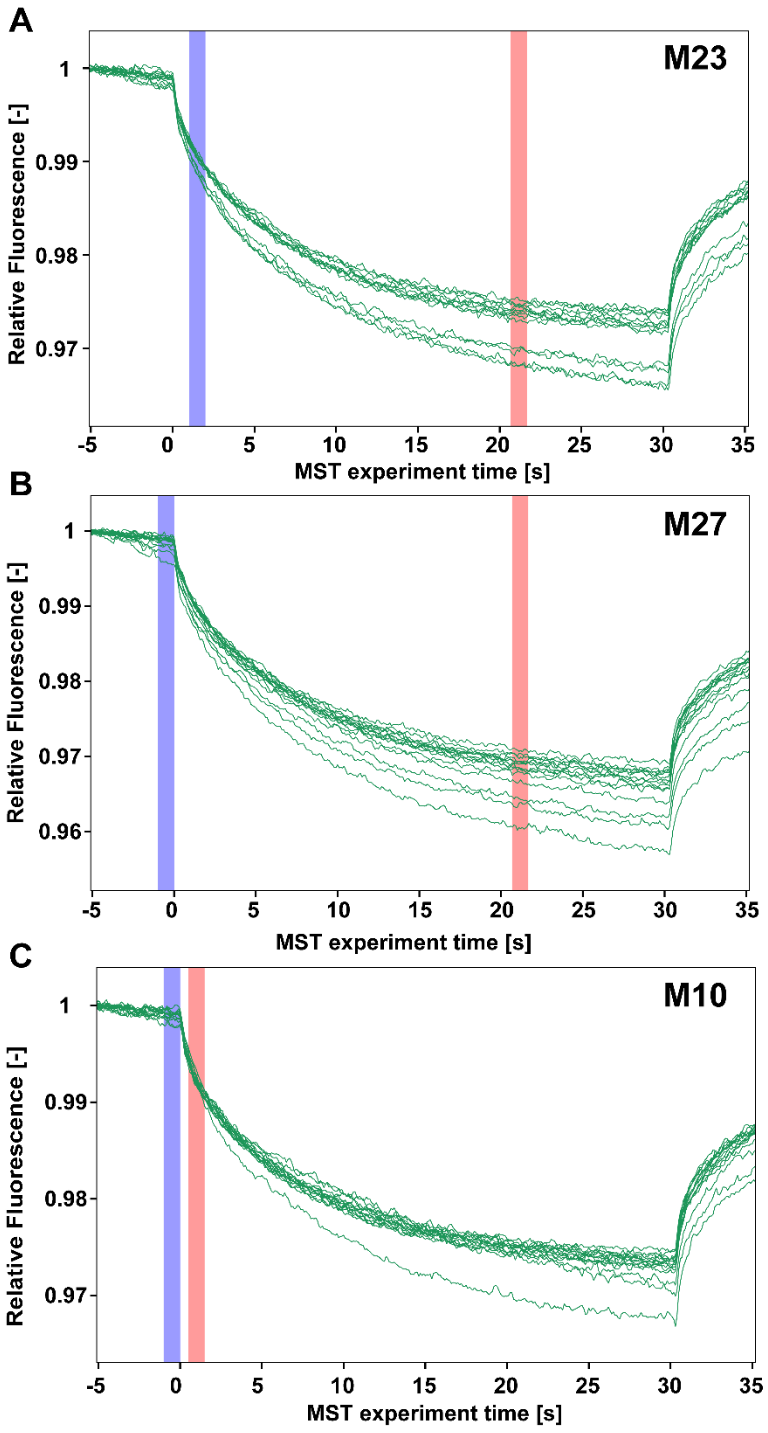


**Figure S4. MST traces of the binding of the inhibitor compounds to NHR2**. Hit compounds, **(A) M23**, **(B) M27**, **(C) M10,** binding to NHR2 detected by MST assay is shown. Titration of compounds to a constant concentration of Alexa488 dye-labeled NHR2 induces a change in thermophoresis. Regions of the traces corresponding to T-jump and thermophoresis signals were used (blue and red bars) for obtaining apparent $K_D$ values, that is, **M23** was analyzed by thermophoresis, **M27** was analyzed by T-jump and thermophoresis, and **M10** was analyzed by T-jump. These methods were chosen based on the response amplitude and/or standard error of the MST traces.

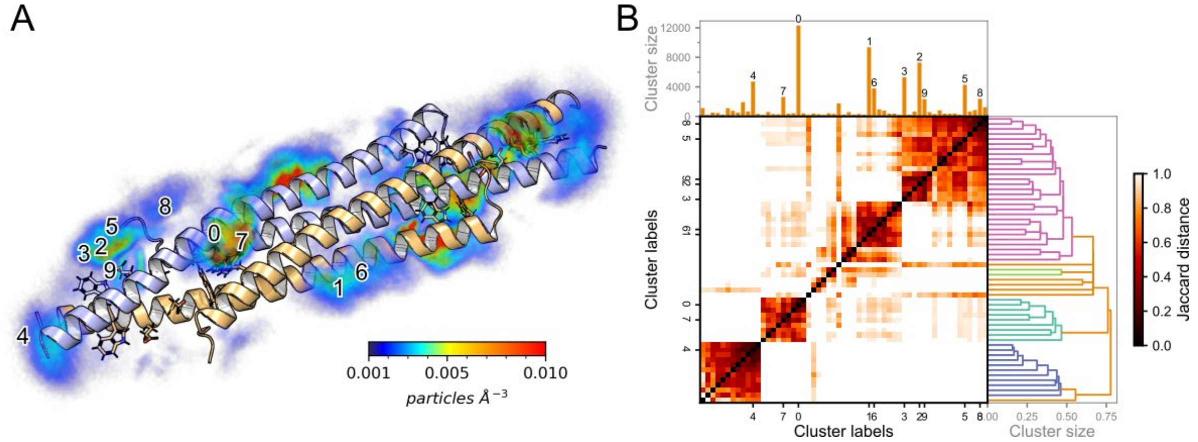


**Figure S5. Binding of M23 to the NHR2 tetramer. (A)** Heavy-atom particle density of **M23** around the NHR2 tetramer is mapped in a color-coded manner (see color scale). A clustering of ligand poses was performed based on interaction fingerprints with NHR2 residues. **(B)** Similarity between the clusters is shown for all clusters with a population > 0.1% of the total number of samples. Clusters with a population > 1.0% are labeled in the order of decreasing population: 0 (largest) to 9 (smallest).

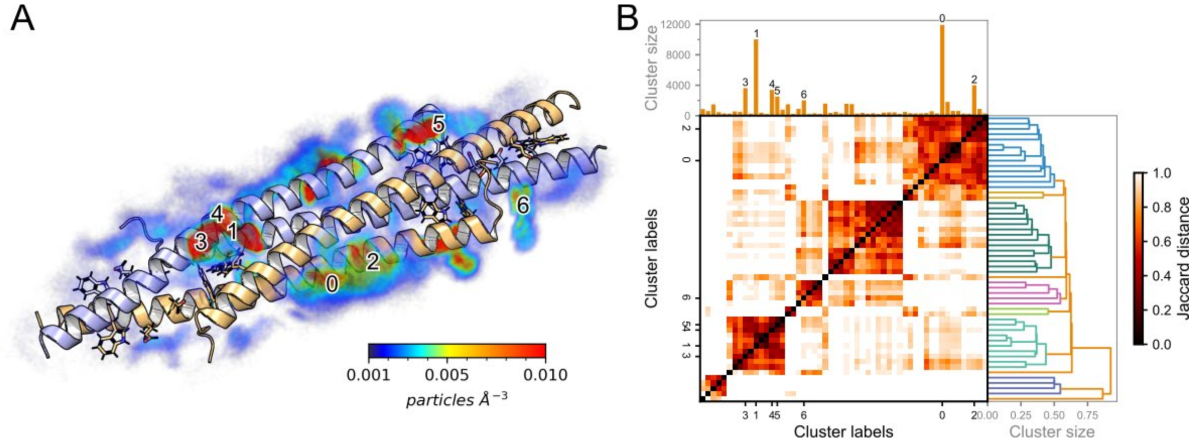


**Figure S6. Binding of M27 to the NHR2 tetramer. (A)** Heavy-atom particle density of **M27** around the NHR2 tetramer is mapped in a color-coded manner (see color scale). A clustering of ligand poses was performed based on interaction fingerprints with NHR2 residues. **(B)** Similarity between the clusters is shown for all clusters with a population > 0.1% of the total number of samples. Clusters with a population > 1.0% are labeled in the order of decreasing population: 0 (largest) to 6 (smallest).

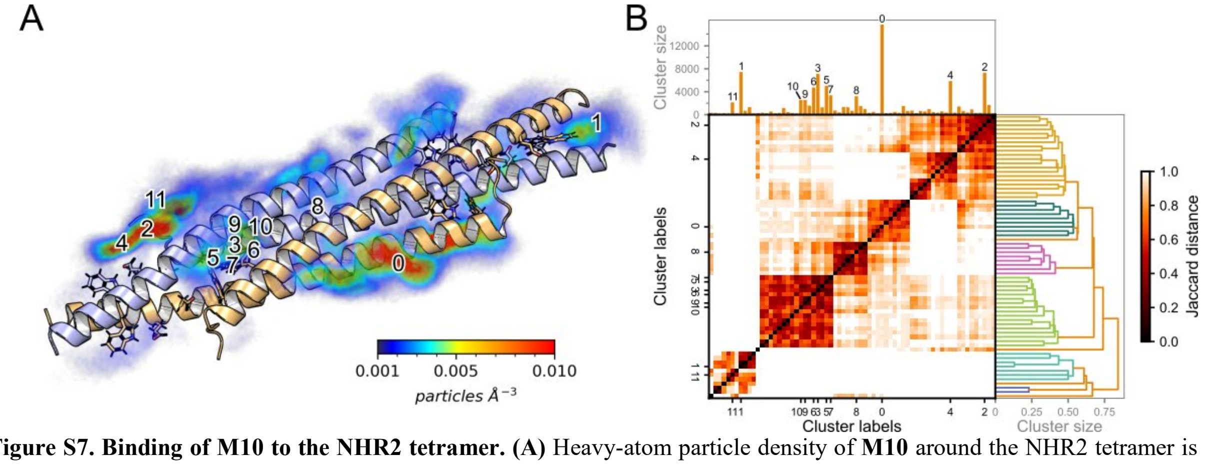


**Figure S7. Binding of M10 to the NHR2 tetramer. (A)** Heavy-atom particle density of **M10** around the NHR2 tetramer is mapped in a color-coded manner (see color scale). A clustering of ligand poses was performed based on interaction fingerprints with NHR2 residues. **(B)** Similarity between the clusters is shown for all clusters with a population > 0.1% of the total number of samples. Clusters with a population > 1.0% are labeled in the order of decreasing population: 0 (largest) to 10 (smallest).

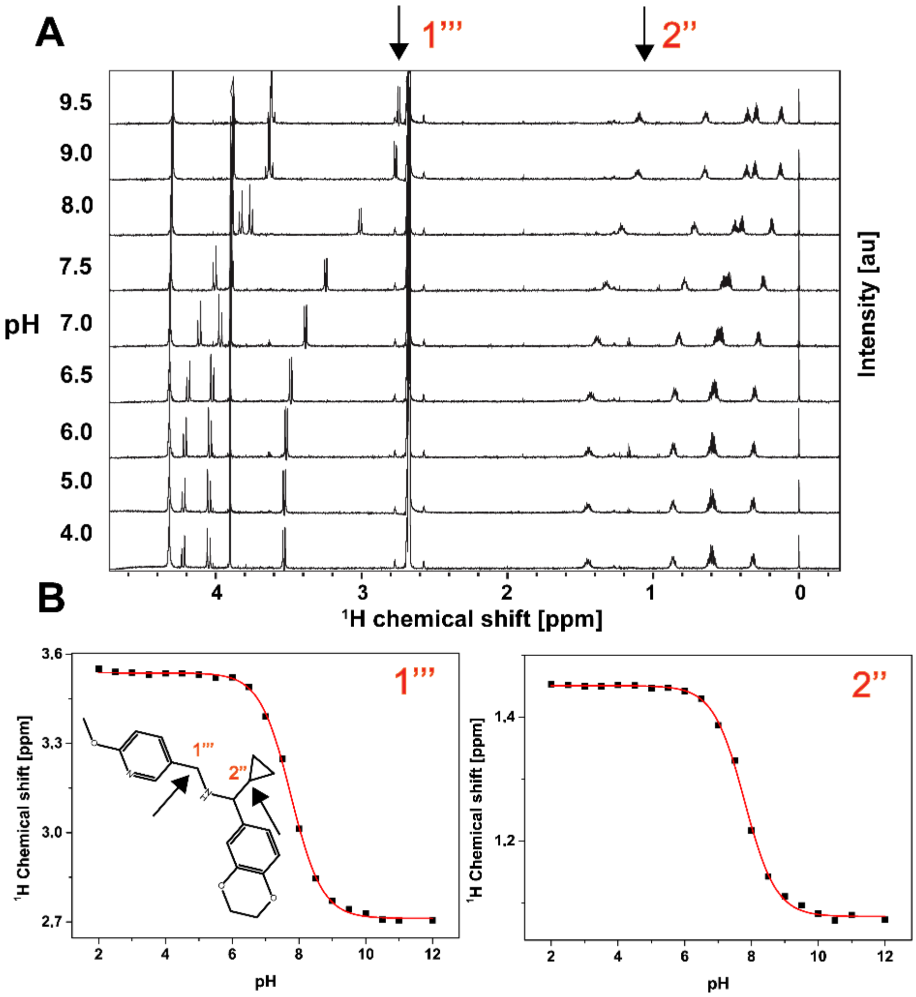


**Figure S8. Determination of the $pK_a$ value of M27 by NMR**. (**A**) Resolved high-field portion of the 700 MHz $^1$H NMR spectrum of 300 µM **M27** measured in the pH range of 2 to 13 in 50 mM sodium phosphate, 100 mM sodium chloride, 10% (v/v) $D_2O$, 10% (v/v) DMSO-$d_6$. Sodium 2,2-dimethyl-2-silapentane-5-sulfonate (DSS) was used for chemical shift referencing and calibrated to a zero ppm value. (**B**) Chemical shift values of reporter protons (labeled as 1''' and 2'', marked by arrows) were plotted against pH. The $pK_a$ value was calculated by fitting to the Henderson-Hasselbalch equation. The $pK_a$ value of the secondary amine is 7.76 ± 0.01 and 7.79 ± 0.02 as indicated by protons 1''' and 2'', respectively.

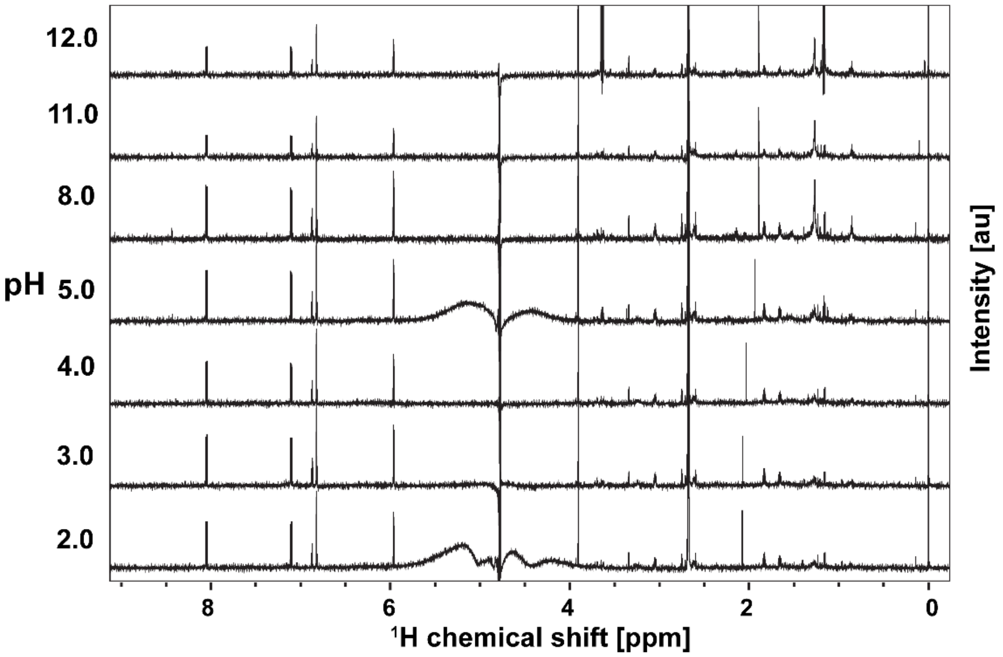


**Figure S9. M10** does not contain a labile proton and did not show a chemical shift change during pH titration.

### Supplementary tables

**Table S1. Molecules selected from ligand-based virtual screening for experimental testing**. Molecules with an $IC_{50}$ value < 1 mM in the cell viability assay and a positive signal in the STD NMR experiment are marked in green.

| Compound ID | ZINC ID[a] | MolPort ID[a] | Company[b] | Catalog ID[b] | MW[c] | xlogP[d] |
|---|---|---|---|---|---|---|
| **M1** | ZINC2698109 | MolPort-000-479-588 | Life Chemicals Inc | F1533-0315 | 292 | 1.12 |
| **M2** | ZINC4279993 | MolPort-000-466-718 | Life Chemicals Inc | F1278-0182 | 322 | 2.51 |
| **M3** | ZINC13628649 | MolPort-000-842-332 | Eximed | EiM07-18223 | 374 | 3.34 |
| **M4** | ZINC4125019 | MolPort-002-125-097 | ChemBridge Corporation | 7731187 | 272 | 2.40 |
| **M5** | ZINC299739741 | MolPort-039-279-427 | Enamine Ltd | Z1603366153 | 350 | 2.53 |
| **M6** | ZINC238012695 | MolPort-039-278-948 | Enamine Ltd | Z929743448 | 329 | 2.71 |
| **M7** | ZINC1397593 | MolPort-001-843-075 | BIONET/Key Organics Ltd | 7J-537S | 302 | 3.28 |
| **M8** | ZINC4107100 | MolPort-002-886-926 | BIONET/Key Organics Ltd | MS-2031 | 330 | 3.08 |
| **M9** | ZINC49882203 | Not available in MolPort database | Sigma Aldrich | PH006100 | 284 | 3.14 |
| **M10** | ZINC1389510 | MolPort-002-869-018 | BIONET/Key Organics Ltd | 4N-031 | 314 | 3.41 |
| **M11** | ZINC2549649 | MolPort-006-755-951 | BIONET/Key Organics Ltd | MS-6536 | 351 | 4.02 |
| **M12** | ZINC13187875 | MolPort-000-183-134 | Enamine Ltd | Z85969754 | 357 | 2.41 |
| **M13** | ZINC8312526 | MolPort-004-185-300 | Enamine Ltd | Z85881800 | 340 | 2.66 |
| **M14** | ZINC13257945 | MolPort-005-386-809 | Enamine Ltd | Z224125632 | 359 | 2.93 |
| **M15** | ZINC8274274 | MolPort-005-517-016 | Enamine Ltd | Z85881817 | 352 | 2.32 |
| **M16** | ZINC42338549 | MolPort-009-327-813 | Enamine Ltd | Z477284616 | 373 | 3.03 |
| **M17** | ZINC97035318 | MolPort-029-929-656 | Enamine Ltd | Z1603203274 | 338 | 3.32 |
| **M18** | ZINC5344491 | MolPort-002-877-145 | BIONET/Key Organics Ltd | 7J-549S | 306 | 3.92 |
| **M19** | ZINC1397637 | MolPort-002-877-148 | BIONET/Key Organics Ltd | 7J-571S | 302 | 4.09 |
| **M20** | ZINC216966461 | MolPort-035-759-722 | ChemBridge Corporation | 16303929 | 333 | 3.10 |
| **M21** | ZINC95384280 | MolPort-028-594-163 | ChemBridge Corporation | 81622110 | 371 | 2.72 |
| **M22** | ZINC32759522 | MolPort-009-382-435 | Enamine Ltd | Z106335344 | 371 | 3.01 |
| **M23** | ZINC9994366 | MolPort-004-217-884 | Enamine Ltd | Z106912176 | 399 | 3.71 |
| **M24** | ZINC58405221 | MolPort-009-590-667 | Enamine Ltd | Z1142473439 | 314 | 2.05 |
| **M25** | ZINC89905076 | MolPort-027-668-844 | Enamine Ltd | Z1492372136 | 323 | 3.20 |
| **M26** | ZINC89762015 | MolPort-028-820-057 | Enamine Ltd | Z152080224 | 403 | 2.21 |
| **M27** | ZINC97034212 | MolPort-029-929-350 | Enamine Ltd | Z1595536589 | 326 | 3.10 |
| **M28** | ZINC170071403 | MolPort-030-014-742 | Enamine Ltd | Z1624129719 | 329 | 3.12 |
| **M29** | ZINC22703670 | MolPort-006-867-651 | Enamine Ltd | Z18362869 | 317 | 1.82 |
| **M30** | ZINC328594644 | MolPort-039-294-016 | Enamine Ltd | Z1839596229 | 343 | 2.73 |

[a] The ZINC ID and MolPort ID of the stereoisomers based on which the compound was chosen for the experimental testing.
[b] All 30 compounds purchased from the vendors are listed along with their catalog numbers at the time of purchase. The actual configuration of the compounds was not redetermined by us. 12 compounds tested in more detail are highlighted in green.
[c], [d] The molecular weight (MW) and xlogP values were taken from the ZINC database.

**Table S2**. Permeability data of reference compounds from Gopalswamy et al. (1).[a]

| **Compound** | **log $P_{eff/PMF}$ $P_0^{-1}$ [b]** | **log $P_{eff/PAMPA}$ $P_0^{-1}$ [c]** |
|---|---|---|
| progesterone | 0.70 ± 0.28 | -4.94 |
| theophylline | 0.12 ± 0.50 | -5.91 |
| pralidoxime (P2-PAM) | -9.71 ± 0.74 | -7.52 |
| atropine | -0.75 ± 0.94 | -5.26 |
| chlorpromazine | 1.58 ± 0.18 | -5.26 |
| diazepam | 1.10 ± 0.16 | -5.40 |
| asoxime (HI-6) | -14.43 ± 0.91 | -7.69 |
| methoxime (MMB4) | -10.67 ± 1.32 | -9.25 |
| promazine | 1.46 ± 0.17 | -4.88 |

[a] Note that for some of the values, typos have been corrected here compared to the respective table in ref. (1). All computations in ref. (1) were done with the correct values.

[b] $P_{eff/PMF}$, effective permeability in cm sec$^{-1}$ calculated from free energy calculations (eq. S2). $P_0$, unit factor corresponding to 1 cm sec$^{-1}$. Errors correspond to the standard deviation obtained from calculating the permeability when dividing 50 ns into ten independent 5 ns simulation slices.

[c] $P_{eff/PAMPA}$, effective permeability in cm sec$^{-1}$ obtained from PAMPA assays (2). $P_0$, unit factor corresponding to 1 cm sec$^{-1}$.

### Supplementary methods

#### Permeability estimation from molecular dynamics simulations

##### Molecular dynamics simulations

For setting up molecular dynamics (MD) simulations, atom types and their corresponding parameters were obtained from the AMBER GAFF2 force field (3), using antechamber. The restrained electrostatic potential (RESP) method was used to assign partial charges from electrostatic potentials calculated at the HF/6-31G* level of theory with Gaussian 09 (4). Each simulation system was packed in a 75x75 $Å^2$ 1,2-dioleoyl-*sn*-glycero-3-phosphocholine (DOPC) bilayer plane within a 100 Å tall simulation box using PACKMOL-Memgen (5), using TIP3P water molecules for the water phase (6), and one ligand in the center of the bilayer. $Cl^-$ ions were added to neutralize the system when positively charged compounds were simulated.

All MD simulations were performed using AMBER22 (3). The minimization of the systems was performed using the MPI implementation of PMEMD (7). 5000 cycles of steepest descent were followed in the first step of minimization by conjugate gradient minimization for a total of 10000 cycles. Only water molecules (and ions, if included) were minimized initially, using harmonic restraints of 25 kcal $mol^{-1}$ $A^{-2}$ on the rest of the system. In the second minimization step, the harmonic restraints were decreased to 5 kcal $mol^{-1}$ $A^{-2}$, while in the third and fourth steps of minimization, the restraints were applied only on the ligand. The fifth minimization step was performed without restraints. The minimized systems were then heated from 0 to 100 K for 5 ps in the NVT ensemble. The temperature was controlled using Langevin dynamics with a coupling constant of 1 $ps^{-1}$, SHAKE, and a time step of 2 fs in all cases (8). Further heating to 300 K was performed under NPT conditions, using the Berendsen barostat with semiisotropic pressure scaling along the membrane plane for 115 ps. The simulations were further relaxed under the same conditions until 5 ns were obtained, keeping the compound molecules at the center of the membrane bilayer with a harmonic restraint along the membrane normal (z-axis) of 2.5 kcal $mol^{-1}$ $A^{-2}$.

The permeabilities of the selected compounds were calculated in a similar fashion as done by us and elsewhere (1,9). In brief, steered MD simulations were carried out for each molecule, pulling for 32 ns at 300 K with a force constant of 10 kcal $mol^{-1}$ $A^{-2}$ and a pulling speed of 1 Å $ns^{-1}$ along the membrane normal (z-axis). A total of 33 umbrella windows were extracted, covering 0 to 32 Å with respect to the membrane center along the z-axis. In umbrella sampling simulations, each extracted structure was simulated for 100 ns, maintaining as a reaction coordinate the distance to the membrane center along the z-axis in the umbrella window with a force constant of 2.5 kcal mol$^{-1}$ Å$^{-2}$. The initial 50 ns of each simulation were considered as equilibration time, with the last 50 ns of each simulation being used for further analysis.

**Permeation potential of mean force and permeability calculations**

Potential of mean force (PMF) profiles were calculated from the distance distributions obtained from the umbrella sampling simulations, using the weighted histogram analysis method (WHAM) (10). A value of zero was assigned for the molecule in bulk water (**Figure 8A**). The permeability of the compounds was calculated following the same protocol as done by us before (1). Briefly, as described by Hummer (11), the resistivity ($R$) to the permeation through the membrane of the compounds (eq. S1) can be computed by using the PMF profile ($\Delta G(z)$) and the diffusion along the membrane normal ($z$-axis) ($D(z)$) (12,13):

$$R(z) = \frac{e^{\beta(\Delta G(z))}}{D(z)} \qquad \text{(S1)}$$

where $\beta$ is the inverse of the Boltzmann constant times the absolute temperature, $D\big(z = (z)\big) = \frac{var(z)}{\tau_z}$ and $\tau_z$ corresponds to the characteristic time of the $z$-position autocorrelation in the given window. The inverse of the integral of the resistivity ($R$) along the $z$-axis (eq. S2) results in the effective permeability $P_{\text{eff}}$ (12) (eq. S2):

$$P_{eff} = \frac{1}{R} = \frac{1}{\int_0^z R(z)\, dz} \qquad \text{(S2)}$$

where the range 0 to $z$ covers the width of the whole membrane (2,14). Errors of the calculated permeabilities were estimated by performing the calculations in ten slices of 5 ns from the total 50 ns used for the analysis.

The calibration curve obtained previously (1) for the compounds with experimentally determined PAMPA data of the reference molecules by Bennion et al. (2) (log $P_{\text{eff/PMF}}$ $P_0^{-1}$ and log $P_{\text{eff/PAMPA}}$ $P_0^{-1}$, respectively, **Table S2, Figure 8B**) was used to calculate PAMPA permeabilities for the compounds in **Table 2**.
